# Developmental CA2 perineuronal net reduction restores social memory in *Shank3* mutant mice

**DOI:** 10.1101/2025.10.12.681906

**Authors:** Emma J. Diethorn, Aarushi B Rathaur, Brayan R. Ruiz Lopez, Chino K. Eke, Elizabeth Gould

**Author notes:** Address correspondence to: Elizabeth Gould PhD, Princeton Neuroscience Institute, Princeton University.

## Abstract

Individuals with Autism Spectrum Disorder (ASD) and related neurodevelopmental conditions, like Phelan-McDermid syndrome (PMDS), exhibit social recognition deficits. Previous reports using *Shank3B* knockout (KO) mice, a genetic model with relevance to ASD and PMDS, revealed deficits in social memory in adulthood, but the postnatal onset and mechanisms underlying this dysfunction remain unknown. In the hippocampus, area CA2 contributes to the emergence of social memory during development and remains important for this ability through the lifespan. Perineuronal nets (PNNs), extracellular matrix structures, support social memory in CA2 and help orchestrate critical periods of plasticity in other brain regions. We found that *Shank3B* KO pups exhibit specific deficits in short-term social recognition and social novelty preference, which become evident at the end of the second postnatal week and persist into adulthood along with CA2 network aberrations. Excessive PNNs in area CA2 were detected in KO pups at the postnatal time when social recognition function typically emerges in healthy pups. This time was also characterized by greater sequestration of the guidance cue semaphorin-3A (sema3A) and overgrowth of afferents from the supramammillary nucleus (SuM) in KOs, suggesting that inordinate PNNs during a sensitive period may disorganize developing CA2 circuitry. Reduction of CA2 PNN levels prior to the onset of social dysfunction recovered social recognition function during development, as well as reduced sema3A and SuM inputs to the CA2. The restoration of behavioral function persisted into adulthood along with partial normalization of CA2 network activity. Together, these findings reveal how excess PNN formation in the developing hippocampus may give rise to impaired social memory by disrupting afferent input, effects that are reversible by early life intervention.

## Introduction

Autism Spectrum Disorder (ASD) is known to have a multifactorial etiology with both environmental and genetic factors implicated. In genome-wide association studies, mutations of the SHANK3 gene have been identified as one of several genetic risk factors for ASD (Gauthier et al., 2009; Connolly et al., 2017). In children, deletion or mutation of the SHANK3 gene can contribute to social and cognitive developmental delays that are classified as Phelan-McDermid syndrome (PMDS) (Phelan and McDermid, 2011). Most children with PMDS meet the criteria for ASD diagnosis and have diminished social interest and social novelty preference, in contrast to typically developing children (Weigelt et al., 2012; Guillory et al., 2021). In adult mice with mutations in the *Shank3* gene, including *Shank3B* knockouts (KOs), decreased interest in social stimuli and impaired social recognition have been reported (Tao et al., 2022; Cope et al., 2023). Since this mutation reflects some aspects of neurodevelopmental disorders in humans, characterizing the postnatal onset of social dysfunction is an important first step in investigating the underlying neural mechanisms.

Healthy developing mice display evidence of social recognition function in the first postnatal week of life, which manifests as a physical and vocal preference for the caregiving mom over a novel dam (Laham et al., 2021). As mice develop, their social repertoire expands and preference for novel peers over familiar peers and littermates emerges toward the end of the second postnatal week (Diethorn and Gould, 2023a, b). Social recognition and social novelty preference have been linked to the CA2 region of the hippocampus in adulthood (Hitti and Siegelbaum, 2014; Stevenson and Caldwell, 2014), and specific patterns of network activity, including phase-amplitude coupling and sharp-wave ripples, play important roles in these functions (Oliva et al., 2020; Zhu et al., 2023; Laham et al., 2024). The involvement of area CA2 in social recognition begins during the early postnatal period (Laham et al., 2021; Diethorn and Gould, 2023a) and expands as the region undergoes substantial remodeling characterized by afferent ingrowth and the appearance of perineuronal nets (PNNs) (Dominguez et al., 2019; Carstens et al., 2016; Diethorn and Gould, 2023a, b).

PNNs are extracellular matrix structures that in development have been associated with critical period closure, and in adulthood are thought to modulate plasticity (Wang and Fawcett, 2012; Foscarin et al., 2017; Mirzadeh et al., 2019). Emergence of PNNs in area CA2 during the second postnatal week coincides with the onset of social novelty preference in typically developing mice (Diethorn and Gould, 2023a, b). As mice mature, healthy PNN levels in CA2 continue to support social recognition abilities (Cope et al., 2022; Rey et al., 2022; Alexander et al., 2025; Mehak et al., 2025). In the developing hippocampus and cortex, semaphorin-3A (sema3A) is sequestered by PNNs and influences axon ingrowth and synapse elimination (De Wit et al., 2005; Morita et al., 2006; Yamashita et al., 2007; Vo et al., 2013; Uesaka et al., 2014; Fawcett et al., 2019; Mei et al., 2025). The CA2 region integrates afferent input from many regions that are important for social novelty detection, including the supramammillary nucleus of the hypothalamus (SuM) (Chen et al., 2020; Robert et al., 2021; Li et al., 2023), a pathway known to undergo considerable postnatal remodeling (Diethorn and Gould, 2023a). These findings raise the possibility that social dysfunction in *Shank3B* KO mice arises from atypical expression of CA2 PNNs, which disrupts sema3A signaling and SuM-CA2 circuit development.

To better understand mechanisms of developmental social dysfunction, we characterized social behavior in developing *Shank3B* KO mice and observed deficits in the ability to form a short-term memory of a recently encountered peer in the second postnatal week of life, which persists into adulthood and is accompanied by disrupted CA2 network responses to social stimuli. We also observed that KO pups exhibit elevated PNNs in CA2, along with excessive sema3A and SuM ingrowth, when social dysfunction is first apparent. Lastly, we found that lowering CA2 PNNs in KOs during the second postnatal week reduces developmental sema3A and SuM ingrowth, while restoring social function. Restoration of social recognition function and some aspects of CA2 network activity were evident in adult KOs after developmental reduction of PNNs. Together, these findings suggest that atypical CA2 PNNs contribute to impaired social recognition and network activity in *Shank3B* KO mice by disrupting postnatal development of CA2 circuitry, and that reduction of PNNs during development allows for healthy CA2 function to emerge and persist into adulthood.

## Methods

### Animals

All mice were cared for in accordance with the Princeton University Institutional Animal Care and Use Committee. Mice were group-housed and kept on a 12:12hr light/dark cycle in Optimice cages with food and water received ad libitum. Mixed groups of male and female mice were used for all experiments. Male and female *Shank3B* heterozygous mice were purchased from Jackson Laboratories (strain #017688) and paired together for breeding, yielding wildtype (WT), knockout (KO), and heterozygous (HET) offspring. Pups were genotyped (Transnetyx) via ear punch prior to P14 and received individual toe tattoos for identification. Pups were weaned on P21 and group-housed by sex.

### Behavioral Testing

#### Ultrasonic Vocalization Recording

Ultrasonic vocalizations (USVs) were recorded from P5 KO and WT pups in isolation to determine whether basic vocalization would be typical in pups of this model. P5 pups were placed individually in a 12”x12” matte plexiglass testing arena with a white base and black walls, maintained at 32°C to simulate nest temperature, and USVs were recorded for 2min. An Avisoft-UltraSoundGate 116Hb kit with a single microphone and Avisoft RECORDER software (v4.2.29) were used with the following parameters adapted from Laham et al., 2021: 300kHz sampling rate, 16-bit format, 0.032s buffer, 50% overlap and 256 FFT size. Sound files were processed using Avisoft SASLab Pro (v5.2.13) and put through a high-pass filter and an average image filter to exclude frequency under 30kHz and background noise, respectively. Spectrograms were then created with the following parameters: 512 FFT size, 100% frame rate, Hamming window, 75% temporal overlap. USVs were detected with automatic whistle tracking using a 2ms minimum duration and 15ms hold time. Calls with a peak amplitude less than -62dB were rejected. From this data, call number, duration (ms), amplitude (dB), and frequency (kHz) were analyzed.

#### Direct Social Interaction Test

Mice were subjected to a direct social interaction test (DSIT) on P14, P21, and in adulthood as described (Diethorn and Gould, 2023a) in the same arena as above. Mice were habituated to the room for 15min prior to behavior, and subject mice were habituated to the arena for 5min before testing. On the day of testing, mice were paired with a novel sex- and age-matched (± 2 PND) HET stimulus mouse (or a familiar HET littermate for the P14 littermate DSIT) and allowed to interact freely for 5min. After an inter-trial interval (ITI) during which the mice were returned to the home cage, the test mouse and now-familiar stimulus mouse (or a new novel peer for the P14 littermate DSIT) were paired and allowed to interact for an additional 5min. A 1hr ITI was used for P14 and P21 mice to restrain testing to a single postnatal day, while a 24hr ITI was used for adult mice. Both trials were videorecorded, and the amount of time the test mouse spent interacting with the stimulus mouse was manually scored. The apparatus was thoroughly cleaned with 70% ethanol between each mouse and trial.

#### Object Location Test

To assess hippocampus-dependent place memory, mice were subjected to the object location task on P21 and P60 as previously described (Cope et al., 2023). P14 pups were not tested on object location memory because at this age, they failed to investigate the object. The apparatus was the same as above, with the addition of a 2in white vertical stripe along the center of one wall to provide an environmental cue. Mice were habituated to the empty apparatus 2x a day for 5min each for 3 days, beginning on P18 for P21 testing. On Day 4, mice underwent the 10min familiarization phase. Two identical objects were placed in the same orientation in adjacent quadrants of the arena, leaving room for the mouse to walk around the entire object, which was taped down. For P21 mice, testing occurred the same day with a 1hr ITI, while adults were tested after a 24hr ITI following the familiarization phase. For the test, the two identical objects remained the same, but one was moved to an adjacent quadrant and rotated 180°. The moved object was counterbalanced across trials. Both the familiarization and testing trials were recorded and investigation time of the moved vs. unmoved object in the test phase was manually quantified. The apparatus and objects were thoroughly cleaned with 70% ethanol between each mouse and trial. Locomotion in the P60 dataset was tracked automatically with ANY-maze (Stoelting) to confirm previously reports that KO mice are hypoactive (Dhamne et al., 2017; Cope et al., 2023). As expected, we found that KOs exhibited decreased locomotion speed and distance (speed: 0.025m/s ± 0.0027; distance: 7.5m ± 0.7973) compared to WT mice (speed: 0.034m/s ± 0.0039; distance: 10.3m ± 1.188).

### Pup surgery

P10 KO and WT pups were anesthetized with 1-3% isoflurane and transferred to a stereotaxic apparatus with a nose cone adaptor for pups (Kopf 940, Kopf 934-B), padded ear bars, and an infrared heating pad (Kent Scientific). Pups received subcutaneous injections of saline to hydrate, and of Carprofen for pain relief (5mg/kg). The scalp was incised and the head was leveled using bregma and lambda landmarks. Dorsal CA2 was targeted bilaterally at the following stereotaxic coordinates: AP= -1.80mm, ML= ±1.60mm, DV= -1.60mm. Chondroitinase ABC from Proteus vulgaris (chABC) (15nL, 18U/mL) (Sigma-Aldrich, C2905-10UN) or penicillinase (pnase) (15nL, 18U/mL) (Sigma-Aldrich, C0389) were prepared in 0.1% BSA in saline, loaded into a 10μl Nanofil syringe (WPI) fitted with a 33g beveled needle (WPI), and infused into CA2 at a rate of 5nL/min. This volume (15nL) was chosen based on cresyl violet injections indicating little to no spread beyond CA2. The scalp was then closed with nylon sutures (Ethilon).

### Electrophysiology

#### Electrode Implantation

To assess CA2 network activity in KOs with social recognition dysfunction, young adult mice were implanted with an electrode in the CA2 region. Although there was a robust statistically significant social recognition deficit in adult KOs compared to WTs, we observed considerable variability in both groups in social preference (Fig. 1e,f). To avoid recording from mice that were outliers in either group, a pre-implant DSIT was used to select for implantation WTs (or chABC-treated KOs) with a discrimination index >0.24, indicating a strong novelty preference, and KOs (or pnase-treated KOs) with an index <0.08, indicating no social preference. Adult mice were implanted with a custom-made unilateral electrode implant with 4 platinum/iridium probes spaced 100μm apart (MicroProbes) into dorsal CA2 of the right hemisphere. Mice were mounted into the stereotaxic apparatus and prepared as described above. Self-tapping bone screws (FST, #19010-10) were implanted on either side of the midline and anterior to bregma for stabilization. A ground screw was inserted contralateral to the electrode, wrapped with silver ground wire, and covered with conductive paint (MG Chemicals, #842AR). The electrode was then implanted at the following coordinates: AP= -1.82mm, ML= +2.15mm, DV= -1.67mm. C&B-Metabond dental cement (Parkell, S280) was used to stabilize the headstage to the skull and ground screw, and a second cement (Keystone, #0921090) was applied to further encase the implant. A threaded base of a 3D-printed protective cap (available at https://github.com/diethorn/3DPrinting) was embedded into the cement surrounding the headstage, and the top of the cap was screwed onto the base to protect the input port.

**Figure 1:**
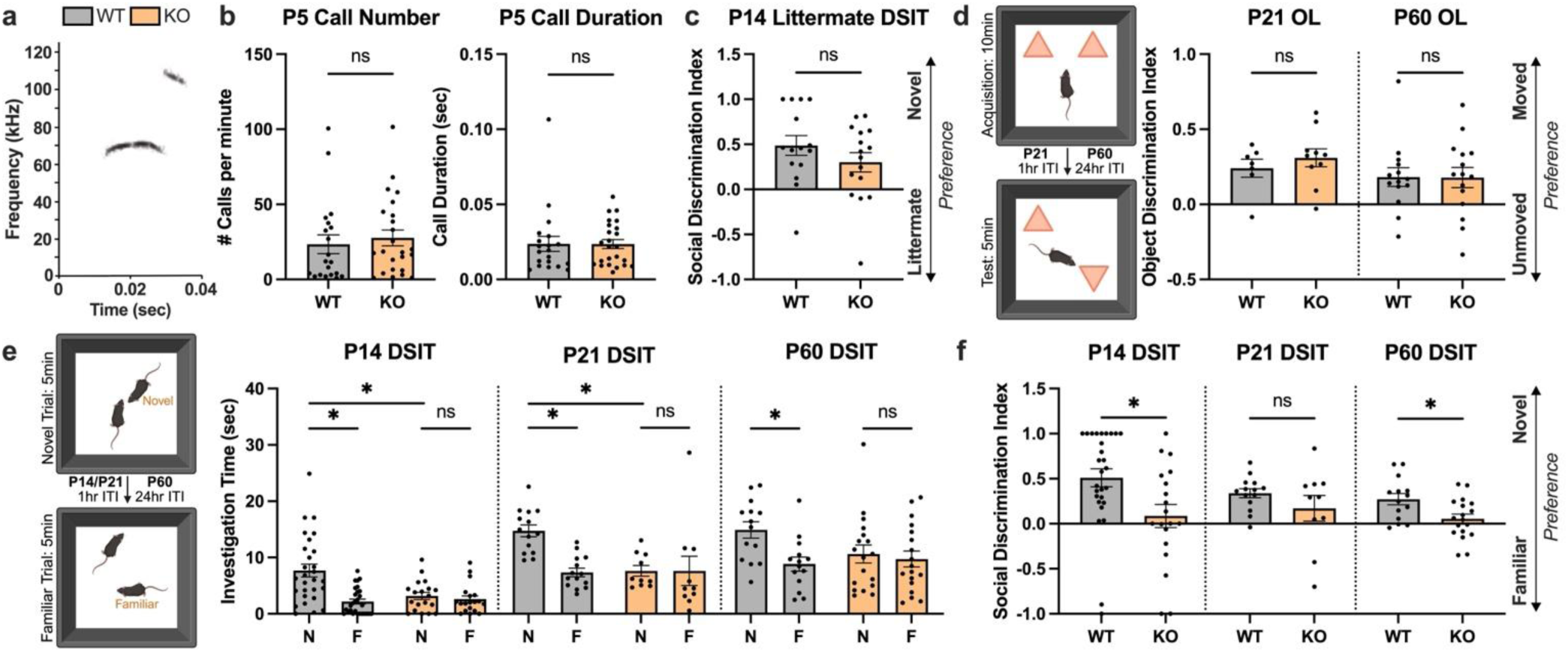
*Shank3B* KO mice demonstrate partially impaired social recognition during development and in adulthood but exhibit some typical social and nonsocial behavior. **a)** Representative USV spectrogram from a P5 KO mouse. **b)** No differences were seen between genotypes in isolation call number (t_42_ = 0.5225, p = 0.6041) or duration (Mann-Whitney U = 213, p = 0.5360) (WT n = 20, KO n = 24). **c)** In a version of the DSIT, P14 mice were first exposed to a familiar littermate followed by a novel peer. There were no differences between genotypes – KOs and WTs had a positive discrimination index (t_29_ = 1.226, p = 0.2301; WT n = 15, KO n = 16). **d)** P21 and young adult mice were subjected to an object location task to assess nonsocial hippocampus-dependent memory, and both genotypes at P21 (t_15_ = 0.7940, p = 0.4396; WT n = 7, KO n = 10) and P60 (t_27_ = 0.0356, p = 0.9719; WT n = 14, KO n = 15) similarly preferred the moved object. **e)** In the DSIT with a novel and then familiar peer, P14 WT pups exhibited preference for social novelty while KO pups did not and, additionally, they investigated the novel peer less than WTs (interaction: F_(1, 45)_ = 11.21, p = 0.0016; WT: p < 0.0001, n = 28; KO: p = 0.8516, n = 19; novel: p = 0.0004). The same effects were seen in the P21 DSIT (interaction: F_(1, 22)_ = 6.899, p = 0.0154; WT: p = 0.0010, n = 14; KO: p > 0.9999, n = 10; novel: p = 0.0015). In adulthood, WTs exhibited a strong preference for the novel peer, while KOs again failed to do so (interaction: F_(1, 30)_ = 9.212, p = 0.0049; WT: p < 0.0001, N = 14; KO: p = 0.6845, n = 18). **f)** The social discrimination index was significantly lower in P14 KO mice compared to WTs (t_45_ = 2.633, p = 0.0115). Although this measure was not statistically different between genotypes on P21 (Mann-Whitney U = 54, p = 0.3712), it was at P60 (t_30_ = 2.650, p = 0.0127), indicating persistent dysfunction in social memory capabilities. * p<0.05; ns, not significant.

#### Electrophysiological Recording

Local field potential was recorded in tethered, freely moving mice from up to 4 electrodes placed in the CA2 using an Omniplex recording system (Plexon, Inc.) at a sampling rate of 40kHz. All recordings were conducted in a 12”x12” plexiglass arena in dim lighting, and mice were habituated to the arena prior to the experiment. First, recordings were made over a 3min pre-social baseline period while alone in a familiar arena. Immediately after, recordings were made during a 5min exposure to a novel peer. Post-social recordings were conducted 1hr later for 5min while the animal was alone in its home cage. 24hr after exposure to the novel peer, a final recording was conducted during exposure to the same now-familiar peer for 5min. These recording sessions were chosen to capture times when acquisition, consolidation, and retrieval of social memories were likely occurring. For analysis of LFP data during immobility bouts and social investigation bouts, trials were recorded using a GoPro Hero8 and behavioral bouts were manually detected using BORIS (v7.13.9). Immobile bouts were characterized by lack of walking and included behaviors such as grooming and rearing when not in contact with the stimulus mouse. In the post-social home cage, periods of stationary eating, digging, and nesting were also counted as immobile bouts. Social investigation was defined as periods of face-to-face sniffing, anogenital sniffing, close following, and cooperative rearing in the novel and familiar recording sessions.

#### Electrophysiological Analyses

Custom Python scripts (available at https://github.com/diethorn/ephys) were used to analyze electrophysiological data. For power analyses, LFPs were downsampled to 1kHz, notch-filtered at 60Hz, and band-pass filtered from 1-140Hz. Power spectral density (PSD) was estimated using Welch’s method. Power in theta (4-12Hz) (Colgin, 2016), slow gamma (20-50Hz) (Carr et al., 2012), and fast gamma (50-140Hz) (Balakrishnan and Pearce, 2015) ranges was quantified by integrating the area under the PSD curve. Aperiodic (1/f^x^) component analyses of the exponent and offset (i.e. slope and intercept) were conducted according to Donoghue et al., 2020. PSDs were fit within 2-40Hz using the SpecParam toolbox with a knee-mode model, and the aperiodic components were extracted from these fits. To detect sharp-wave ripples (SWRs), a third-order Butterworth band-pass filter was applied to isolate 140–220Hz. The filtered signal was Hilbert-transformed to compute the amplitude envelope, followed by Gaussian smoothing. The smoothed envelope was z-scored, and ripple events were identified as periods exceeding 3.5 standard deviations above the mean, ending when the signal dropped below 1.5 standard deviations. SWRs <20ms, >125ms, or within 200ms of another were excluded. Additionally, SWRs with mean amplitudes >0.4mV were considered outliers and removed. SWR rate during behavioral bouts was determined by dividing the number of ripples by total bout duration. Phase-amplitude coupling (PAC) between theta and slow gamma was quantified using the modulation index (MI) method described by Tort et al., 2010.

### Histology

Mice were perfused on P14, P21, or in adulthood for histological analysis of the CA2 region. Mice were anesthetized with pentobarbital and perfused with cold 4% paraformaldehyde (PFA). Brains were extracted and post-fixed in 4% PFA for 48hr, cryoprotected in 30% sucrose, and stored at -80°C until sectioning. Brains were cut on a Leica 3050S cryostat at 40μm and stored in well plates filled with 0.1M PBS with 0.1% sodium azide. Free-floating immunohistochemistry was performed on P14, P21, and adult tissue. At least 3 sections per brain were stained for each marker.

Sections were first washed in 0.1M PBS for 5min and then transferred to a blocking solution of 3% normal donkey serum (NDS) in 0.1M PBS with 0.3% Triton X-100 for 1.5hr on a shaker. Sections were then incubated in blocking solution plus primary antibodies overnight at 4°C. For PNN labeling, sections were incubated in *Wisteria floribunda agglutinin* (WFA) and rabbit anti-aggrecan (ACAN) or goat anti-ACAN. To examine PV+ interneurons, tissue was labeled with mouse anti-PV. For DG-CA2 afferents, sections were incubated in rabbit anti-zinc transporter 3 (ZnT3). For SuM-CA2 afferents, sections were incubated in rabbit anti-vGlut2 or rat anti-Substance P (SP), as well as rabbit anti-neurokinin 1 receptor (NK1R), which are the active receptors of SP. All reactions contained either mouse anti-RGS14 or rabbit anti-PCP4 to label the CA2 region. Additionally, rabbit anti-c-fos was used for SuM and CA2 IEG labeling. After 24hr, sections were washed in 0.1M PBS for 5min and then transferred to the secondary antibody solution in 0.1M PBS with 0.3% Triton X-100 for 1.5hr. Following secondary antibody incubation, sections were incubated in the counterstain Hoechst 33342 (1:5000) (Molecular Probes, H3570) for 10min and then transferred to 0.1M PBS. Sections were then mounted on Superfrost Plus slides (Fisher Scientific) and left to dry overnight, followed by coverslipping with Vectashield antifade mounting media (Vector Labs). To confirm the placement of each electrode in CA2 (stratum radiatum, pyramidal, or oriens), tissue was labeled with RGS14. Data from any electrode not within a CA2 layer were removed from all analyses. All histochemistry and immunolabeling information can be found in Table 1.

**Table 1:** Reagents and antibodies.

| Primary reagent or antibody | Concentration | Source | Catalog | RRID |
| --- | --- | --- | --- | --- |
| Rabbit anti-cellular FBJ osteosarcoma (c-fos) | 1:500µl | Cell Signaling Technology | 2250 | AB_2247211 |
| Mouse anti-regulator of G-protein signaling 14 (RGS14) | 1:500µl | Antibodies Incorporated | 75-170 | AB_2179931 |
| *Wisteria floribunda* agglutinin (WFA) | 1:1000µl | Sigma-Aldrich | L-1516 | AB_2620171 |
| Rabbit anti-aggrecan (ACAN) | 1:1000µl | Millipore | AB1031 | AB_90460 |
| Goat anti-aggrecan (ACAN) | 1:1000µl | R and D Systems | AF1220 | AB_2219833 |
| Mouse anti-parvalbumin (PV) | 1:500µl | Sigma-Aldrich | P3088 | AB_477329 |
| Rabbit anti-Purkinje cell protein 4 (PCP4) | 1:500µl | Atlas Antibodies | HPA005792 | AB_1855086 |
| Rabbit anti-semaphorin-3A (sema3A) | 1:200µl | Abcam | ab23393 | AB_447408 |
| Rabbit anti-vesicular glutamate transporter 2 (vGlut2) | 1:500µl | Synaptic Systems | 135 403 | AB_887883 |
| Rat anti-Substance P (SP) | 1:500µl | Millipore | MAB356 | AB_94639 |
| Rabbit anti-neurokinin 1 receptor (NK1R) | 1:500µl | Millipore | AB5060 | AB_2200636 |
| Rabbit anti-zinc transporter 3 (ZnT3) | 1:500µl | Alomone Labs | AZT-013 | AB_2756813 |

| Secondary reagent or antibody | Concentration | Source | Catalog | RRID | Used for |
| --- | --- | --- | --- | --- | --- |
| Streptavidin, Alexa Fluor 488 | 1:1000µl | Invitrogen | S32354 | N/A | WFA |
| Streptavidin, Alexa Fluor 568 | 1:1000µl | Invitrogen | S11223A | N/A | WFA |
| Donkey anti-rabbit, Alexa Fluor 488 | 1:500µl (1:200*) | Invitrogen | A21206 | AB_2535792 | c-fos, vGlut2, ZnT3, sema3A* |
| Donkey anti-rabbit, Alexa Fluor 568 | 1:500µl | Invitrogen | A10042 | AB_2534017 | NK1R, ACAN |
| Donkey anti-rabbit, Alexa Fluor 647 | 1:500µl | Invitrogen | A-31573 | AB_2536183 | PCP4 |
| Donkey anti-mouse, Alexa Fluor 568 | 1:500µl | Invitrogen | A10037 | AB_2534013 | RGS14, PV |
| Donkey anti-mouse, Alexa Fluor 647 | 1:500µl | Millipore | AP192SA6 | AB_2687879 | RGS14 |
| Donkey anti-rat, Alexa Fluor 488 | 1:500µl | Invitrogen | A-21208 | AB_141709 | SP |
| Donkey anti-goat, Alexa Fluor 568 | 1:1000µl | Invitrogen | A-11057 | AB_2534104 | ACAN |

Because sema3A is deeply embedded in PNNs, pretreatment is required to reveal antigenic sites during immunolabeling. We used previously published methods (Vo et al., 2013) to label sema3A as follows. Sections were first incubated in 0.1M Tris-HCL containing sodium acetate (0.03M) and chABC (0.1U/mL) for 2hr at 37°C to facilitate antibody binding. After, they were washed with 0.2% TBS-Triton and then incubated in a solution containing TBS, methanol (10%), and hydrogen peroxide (0.3%) for 30min. Then, tissue was blocked in 0.2% TBS-Triton containing 5% NDS for 1hr and incubated overnight at room temperature in blocking solution and rabbit anti-sema3A, mouse anti-RGS14, and the lectin WFA. The following day, sections were incubated in 0.2% TBS-Triton with secondary antibodies.

#### Confocal Imaging and Analyses

Slides were imaged with a 1μm z-step and a 40x objective on a Leica SP8 confocal microscope and analyses were conducted using custom macros in Fiji (Image J) (NIH) after being filtered with a rolling ball radius of 50 pixels. For analyzing WFA+ or ACAN+ PNNs, sema3A, and vGlut2, the RGS14+ pyramidal layer (PYR) was traced and the maximum MGV of the z-stack was averaged across sections; SP and NK1R analyses were confined to the oriens (OR) due to the pattern of afferent input (Borhegyi and Leranth, 1997), while ZnT3 was confined to the stratum lucidum (Diethorn and Gould, 2023a). Background intensity was taken from the corpus callosum and subtracted from the MGV for punctate labels like vGlut2 and SP, yielding net intensity. NK1R exists in the developmental corpus callosum (Barbaresi et al., 2017), so background subtraction was not applied. Particle analyses were conducted on SP images to assess the size and presence of boutons (0.15-7.00μm^2^) from thresholded (RenyiEntropy) z-stacks. C-fos+ cells in the CA2 were manually counted in the RGS14+ CA2 PYR and lateral SuM, and atlas traces were overlaid onto confocal images to identify SuM. For PV, each cell in the RGS14+ PYR was traced and intensity was collected as above, in addition to cell area and cell number. Sema3A images in PNN-corrected tissue were collected with different laser settings than in baseline tissue to accommodate lower labeling intensity.

### Statistical Analyses

All statistical analyses were conducted using Graphpad Prism 10. Two- or three-way ANOVAs with Šidák’s multiple comparisons, or unpaired t-tests were used to assess effects across or within genotype, sex, trial, treatment, and/or age, with repeated measures when applicable. Discrimination indices were calculated as the difference between trials divided by the total across trials. When datasets did not meet the assumptions of parametric statistics, the nonparametric Mann-Whitney U test was used. In the baseline electrophysiology experiment, one KO exhibited zero SWRs during investigation of the familiar partner, preventing calculation of some spectral/temporal measures, so mixed-effects analyses were used. Correlation between sema3A and WFA intensity was conducted using simple linear regression.

## Results

### Developing Shank3B KO mice show impaired social recognition of peers but typical behavior on other social and nonsocial tasks

#### P5 *Shank3B* KO pups exhibit typical ultrasonic vocalizations in isolation

The earliest form of social communication in mice includes the emission of ultrasonic vocalizations from pups when they are separated from their mothers, which is thought to signal the mother to retrieve the pup (Okabe et al., 2010). Consistent with previous findings (Balaan et al., 2019), we found that isolation-induced vocalizations did not differ between KO pups and their wildtype (WT) control littermates in number or in duration (number: t_42_ = 0.5225, p = 0.6041; duration: Mann-Whitney U = 213, p = 0.5360) (Fig. 1a, b). Spectral call properties, including frequency and amplitude, also did not differ between genotypes, further suggesting that KO pups exhibit isolation-induced vocalizations consistent with typically developing pups (frequency: t_42_ = 1.737, p = 0.0897; amplitude: t_42_ = 1.431, p = 0.1597) (Fig. S1a, b).

#### *Shank3B* KO pups exhibit impaired recognition after a single short-term, but not continual long-term, presentation of a social stimulus

To determine whether KO pups exhibit basic memory for a littermate with which they were housed since birth, we conducted a version of the direct social interaction task on P14. Pups were first exposed to a littermate and, after a 1hr ITI, were introduced to a novel sex- and age-matched mouse (a “peer”), which typically developing pups should spend more time investigating (Diethorn and Gould, 2023a). Both WTs and KOs exhibited preference for the novel peer, and mice in both groups had a positive social discrimination index in this task, indicating more time spent with the novel social partner over the familiar littermate (t_29_ = 1.226, p = 0.2301) (Fig. 1c). These results suggest that KO pups can recognize a social stimulus after continual long-term presentation in the nest since birth. Next, we investigated whether KO mice would exhibit typical recognition memory after a short-term presentation of a novel social stimulus across three developmental time points: P14, when preference for novel peers first emerges (Diethorn and Gould, 2023a); P21, when pups transition to weanlings and leave the nest (Brust et al., 2015); and P60, when mice reach young adulthood (Brust et al., 2015). Mice were first exposed to a novel sex- and age-matched peer and, after an ITI, were re-exposed to the same now-familiar peer. At P14, we found that *Shank3B* KO pups did not decrease investigation time between exposure to the novel peer and, 1hr later, the same now-familiar peer, while WT controls did (interaction: F_(1, 45)_ = 11.21, p = 0.0016; WT: p < 0.0001; KO: p = 0.8516) (Fig. 1e). In addition to impaired ability to differentiate between the novel and familiar mouse, KO pups spent significantly less time investigating the novel peer in the initial exposure compared to WTs and demonstrated a lower social discrimination index (novel: p = 0.0004; index: t_45_ = 2.633, p = 0.0115) (Fig. 1f).

The same effect was found a week later, on P21, when WTs again decreased investigation time for the familiar peer while KOs were both unable to so and spent less time with the novel mouse in the initial exposure (interaction: F_(1, 22)_ = 6.899, p = 0.0154; WT: p = 0.0010; KO: p > 0.9999; novel: p = 0.0015) (Fig. 1e). However, the social discrimination index, although lower in KO than WT, was no longer statistically different between genotypes. The lack of a significant effect on this measure could be due to high variability in the KO group (Mann-Whitney U = 54, p = 0.3712) (Fig. 1f). A 1hr ITI was used to constrain training and testing to single days for earlier developmental timepoints, but P60 mice underwent testing with 24hr between trials. As adults, KOs were persistently impaired and did not decrease investigation time between exposure to the novel peer on day 1 and the same now-familiar peer on day 2, while WT adults did, in line with previous findings (interaction: F_(1, 30)_ = 9.212, p = 0.0049; WT: p < 0.0001; KO: p = 0.6845) (Fig. 1e) (Cope et al., 2023). At this age, the discrimination index was significantly lower in KOs compared to WTs (t_30_ = 2.650, p = 0.0127) (Fig. 1f).

#### Nonsocial memory remains intact in developing and adult *Shank3B* KO mice

We investigated whether impaired recognition memory and novelty preference in *Shank3B* KO mice extended to hippocampus-dependent place memory, or if dysfunction was specific to social behavior. KO adults have been reported to have intact object location memory (Cope et al., 2023), but younger ages have not been probed. To test this possibility, P21 and P60 mice underwent an object location task in which typically developing mice investigate a moved object more than an object in a consistent location. P14 WT pups did not investigate objects enough to conduct this task and were therefore excluded from this experiment. Both P21 and P60 WT and KO mice preferred the moved object compared to the unmoved object during the test phase, with no significant difference in the object discrimination index between the genotypes at either age (P21: t_15_ = 0.7940, p = 0.4396; P60: t_27_ = 0.0356, p = 0.9719) (Fig. 1d).

### Shank3B KO adults exhibit aberrant CA2 neuronal activation and network activity during social behavior

Since area CA2 is required for social recognition and novelty preference (Hitti and Siegelbaum, 2014; Stevenson and Caldwell, 2014), we next investigated whether baseline and social novelty-induced neuronal activity differ between KO and WT mice using the immediate early gene c-fos as an indirect measure of neuronal activation. We perfused KO and WT mice directly from the home cage to gather baseline c-fos data, or 1hr after exposure to a novel peer in a testing arena (Fig. 2a). No differences were observed in c-fos+ cell density between genotypes in mice perfused from the home cage (p = 0.9852), and both novel-exposed WT and KO mice exhibited a significant increase in c-fos+ cell density compared to their home cage counterparts (interaction: F_(1, 32)_ = 6.627, p = 0.0149; WT: p < 0.0001; KO: p < 0.0001). However, novel-exposed KOs had significantly lower c-fos+ cell density in CA2 compared to novel-exposed WT mice (p = 0.0022) (Fig. 2b, c). Because c-fos labeling is an indirect marker of neuronal activation that lacks temporal specificity, we next used in vivo electrophysiology of awake behaving mice to examine CA2 network activity between genotypes.

**Figure 2:**
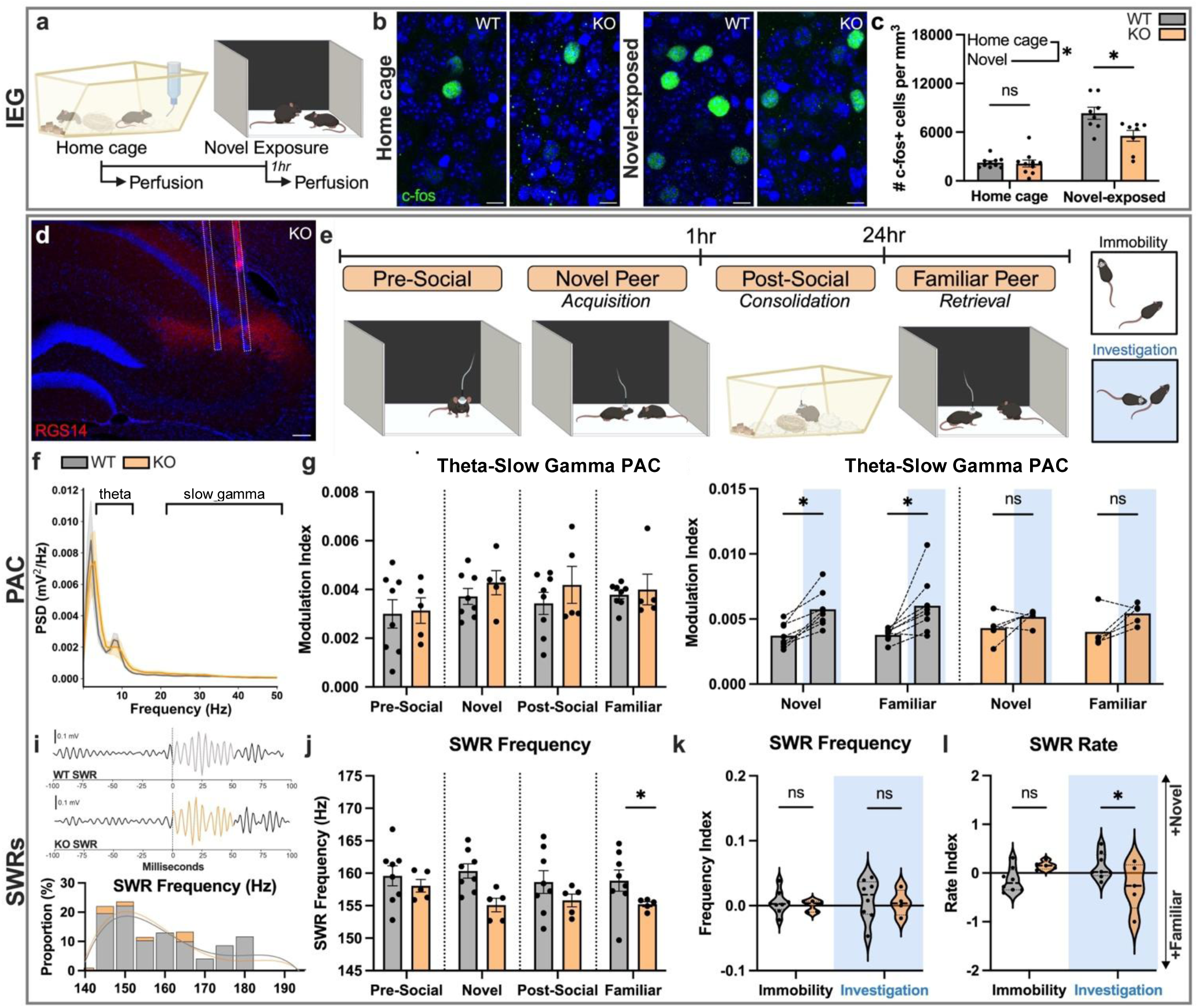
CA2 IEG expression and network activity are atypical in *Shank3B* KO adults. **a)** Schematic of IEG experimental paradigm. **b)** Representative images of c-fos+ cells in the CA2 of adult WT and KO mice perfused from the home cage and 1hr after exposure to a novel peer, counterstained with Hoechst (blue). Scale bars = 50μm. **c)** Mice perfused from the home cage did not differ in CA2 c-fos+ cell density between genotypes (p = 0.9852, n = 10 per genotype). In those exposed to a novel peer, c-fos+ cell density was significantly increased in both genotypes compared to home cage counterparts (interaction: F_(1, 32)_ = 6.627, p = 0.0149; WT p < 0.0001; KO p < 0.0001; n = 8 per genotype). However, novel-exposed KOs had significantly lower c-fos+ cell density than novel-exposed WTs (p = 0.0022). **d)** Representative image of an electrode implant in the RGS14+ CA2 of an adult KO mouse. Scale bar = 100μm. **e)** Schematic of electrophysiological recording paradigm, with the test mouse denoted by the headstage. **f)** PSD plot from representative mice highlighting theta (4-12Hz) and slow gamma (20-50Hz). **g)** Theta-slow gamma PAC is not different between genotypes within recording sessions while immobile (pre-social: t_11_ = 0.1560, p = 0.8789; novel: t_11_ = 0.9917, p = 0.3427; post-social: t_11_ = 0.9198, p = 0.3774; familiar: Mann-Whitney U = 15, p = 0.5237; WT n = 8, KO n = 5). **h)** In WTs, theta-slow gamma PAC was higher during active bouts of social investigation with the novel or familiar peer compared to bouts of immobility within the same trial (bout type: F_(1, 7)_ = 25.06, p = 0.0016; novel: p = 0.0295; familiar: p = 0.0188). This effect was absent in KO mice (F_(1, 4)_ = 5.545, p = 0.0781). **i)** Representative SWR trace and frequency distribution histogram during the novel and familiar trials. **j)** SWR frequency did not differ between WTs and KOs during pre-and post-social sessions but was lower in KOs while with the novel or familiar peer during periods of immobility (pre-social: t_11_ = 0.7228, p = 0.4849; novel: t_11_ = 3.230, p = 0.0080; post-social: t_11_ = 1.200, p = 0.2555; familiar: Mann-Whitney U = 5, p = 0.0295). **k)** No effects were seen in SWR frequency as index scores in either behavioral bout (interaction: F_(1, 10)_ = 0.00005; p = 0.9945). **l)** The SWR rate index was lower in KOs compared to WTs but only during active social investigation (interaction: F_(1, 11)_ = 9.390, p = 0.0108; immobility: p = 0.1738; investigation: p = 0.0478). * p<0.05; ns, not significant.

In CA2, synchronized network events like PAC and SWRs contribute to social recognition function (Oliva et al., 2020; Laham et al., 2024). To assess whether KO mice exhibit aberrations in these oscillations, we recorded from CA2 in the following conditions: 1) a pre-social baseline session; 2) a session with a novel mouse; 3) 1hr later in the home cage, presumably during post-social consolidation; and 4) 24hr after exposure to the novel mouse, with the same now-familiar mouse (Fig. 2d, e). We first checked whether the power spectral densities (PSDs) of KO and WT mice differed in their aperiodic (1/f^x^) components, which represent non-oscillatory neuronal firing in the LFP and are known to be affected by age and disease (Donoghue et al., 2020; Pochinok et al., 2024). Upon finding no differences between genotypes in the aperiodic component’s exponent or offset (i.e. the slope and intercept of the fit) and only moderate fluctuations across recording sessions in both genotypes (Fig. S2a-d), we proceeded with analyzing power, PAC, and SWRs.

Across recording sessions, power was generally comparable between KO and WT mice during periods of immobility (theta: F_(1, 11)_ = 2.675, p = 0.1302; slow gamma: F_(1, 11)_ = 0.3833, p = 0.5484; fast gamma: F_(1, 11)_ = 0.0982, p = 0.7599) (Fig. 2f), and overall gamma power seemed to rise during post-social memory consolidation in both genotypes (slow gamma: F_(1.928, 21.21)_ = 6.250, p = 0.0078, E = 0.9640; fast gamma: F_(1.565, 17.21)_ = 23.11, p < 0.0001, E = 0.7823) (Fig. S2e-g). We found no differences in power between genotypes during active investigation of the novel peer (F_(1, 11)_ = 0.0227, p = 0.8830) but, with the familiar peer on day 2, power was affected in KOs specifically in the slow gamma range (20-50Hz) (F_(1, 11)_ = 6.085, p = 0.0313; slow gamma: p = 0.0154) (Fig. S2h). This effect was not apparent in theta (4-12Hz) (p = 0.0683) nor fast gamma (50-140Hz) (p = 0.6907) power with the familiar peer, although the former approached significance. Given that PAC between theta and gamma within CA2 has been implicated in social recognition function (Zhu et al., 2023; Laham et al., 2024), and because slow gamma power specifically was diminished in KOs during investigation of a familiar peer, we next assessed theta-slow gamma coupling within recording sessions. During periods of immobility, KOs exhibited PAC consistent with that of WTs while with the novel and familiar peer, as well as in pre- and-post social recording sessions (Fig. 2g). However, in WTs, PAC was transiently increased during periods of active social investigation of either the novel or familiar peer (F_(1, 7)_ = 25.06, p = 0.0016; novel: p = 0.0295; familiar: p = 0.0188). This comparison was not significant in KOs, suggesting that during active social investigation, synchrony between theta and slow gamma in CA2 is modulated to a lesser degree than in WTs (F_(1, 4)_ = 5.545, p = 0.0781) (Fig. 2h).

We next investigated whether high frequency SWRs (>140Hz), which have been causally linked to social recognition in healthy mice (Oliva et al., 2020), were atypical in adult *Shank3B* KOs. We found that when mice were immobile during exposure to either a novel or familiar peer, KO mice exhibited lower frequency (Hz) in SWRs, an effect not seen when mice were alone in the pre- and post-social recordings and not during active social investigation (pre-social: t_11_ = 0.7228, p = 0.4849; novel: t_11_ = 3.230, p = 0.0080; post-social: t_11_ = 1.200, p = 0.2555; familiar: Mann-Whitney U = 5, p = 0.0295) (Fig. 2i, j) (Fig. S3b). Other SWR properties like rate, duration, and amplitude were similar between WT and KO mice within recording sessions (Fig. S3a, c, e). When assessed as discrimination index scores comparing the difference between novel and familiar social sessions divided by the sum across both social sessions, SWR frequency, duration, and amplitude were consistent between genotypes during both periods of immobility as well as during active investigation (Fig. 2k) (Fig. S3d, f). Lastly, despite no changes in SWR rate between genotypes within sessions, KOs exhibited a lower rate index score than WTs. Notably, this effect was observed during active bouts of social investigation and not when mice were immobile (interaction: F_(1, 11)_ = 9.390, p = 0.0108; immobility: p = 0.1738; investigation: 0.0478) (Fig. 2l). These electrophysiological results collectively indicate that *Shank3B* KO mice exhibit specific differences in CA2 network events that may suggest diminished capacity for differentiation of neuronal oscillations during novel versus familiar social experiences.

### Excess perineuronal nets in CA2 contribute to impaired social memory development and network dysfunction in Shank3B KO mice

To investigate mechanisms underlying developmental social memory dysfunction in *Shank3B* KO mice, we then characterized the emergence of perineuronal nets (PNNs) in area CA2. CA2 PNNs develop around the time when social memory abilities first emerge (Diethorn and Gould, 2023b), raising the possibility that atypical PNN development in CA2 could underlie impaired social behavior in *Shank3B* KO mice.

#### CA2 perineuronal nets have higher intensity in P14 *Shank3B* KO mice compared to WT littermates

To characterize emerging PNNs, we labeled P14 tissue with the plant lectin *Wisteria floribunda agglutinin* (WFA), which binds to chondroitin sulfate (CS)-GAG chains, as well as with aggrecan (ACAN), the main neuronal CS proteoglycan of PNNs. Optical intensity of both WFA+ and ACAN+ PNNs in the CA2 region was significantly higher in P14 KO compared to WT littermates (WFA: t_21_ = 2.717, p = 0.0129; ACAN: t_21_ = 2.764, p = 0.0116) (Fig. 3a, b). In CA2, PNNs surround both pyramidal cells and parvalbumin+ (PV+) inhibitory interneurons. The labeling increase identified in P14 KOs was most likely caused by increased intensity of PNNs surrounding pyramidal cells, since PNN intensity around PV+ interneurons was similar between genotypes (t_19_ = 0.7416, p = 0.4674) as were other PV+ cell characteristics (Fig. S4a-e). A week later, on P21, differences between genotypes in CA2 PNNs observed at P14 were no longer detectable, with both WFA+ and ACAN+ labeling being similar between KOs and WTs (WFA: Mann-Whitney U = 59, p = 0.9487; ACAN: Mann-Whitney U = 58, p = 0.8977) (Fig. 3c, d). Furthermore, no significant genotype effects were observed in CA2 PNN intensity in young adulthood (P60) (WFA: t_12_ = 0.8142, p = 0.4314; ACAN: t_12_ = 0.5419, p = 0.5978) (Fig. 3e, f), despite finding that social dysfunction persists in KOs at that age. Since studies have shown that PNNs may play a role in organizing neural circuitry during development, we next considered whether excessive PNNs during this specific developmental time in KO mice initially and persistently impairs social recognition, despite the eventual PNN normalization.

**Figure 3:**
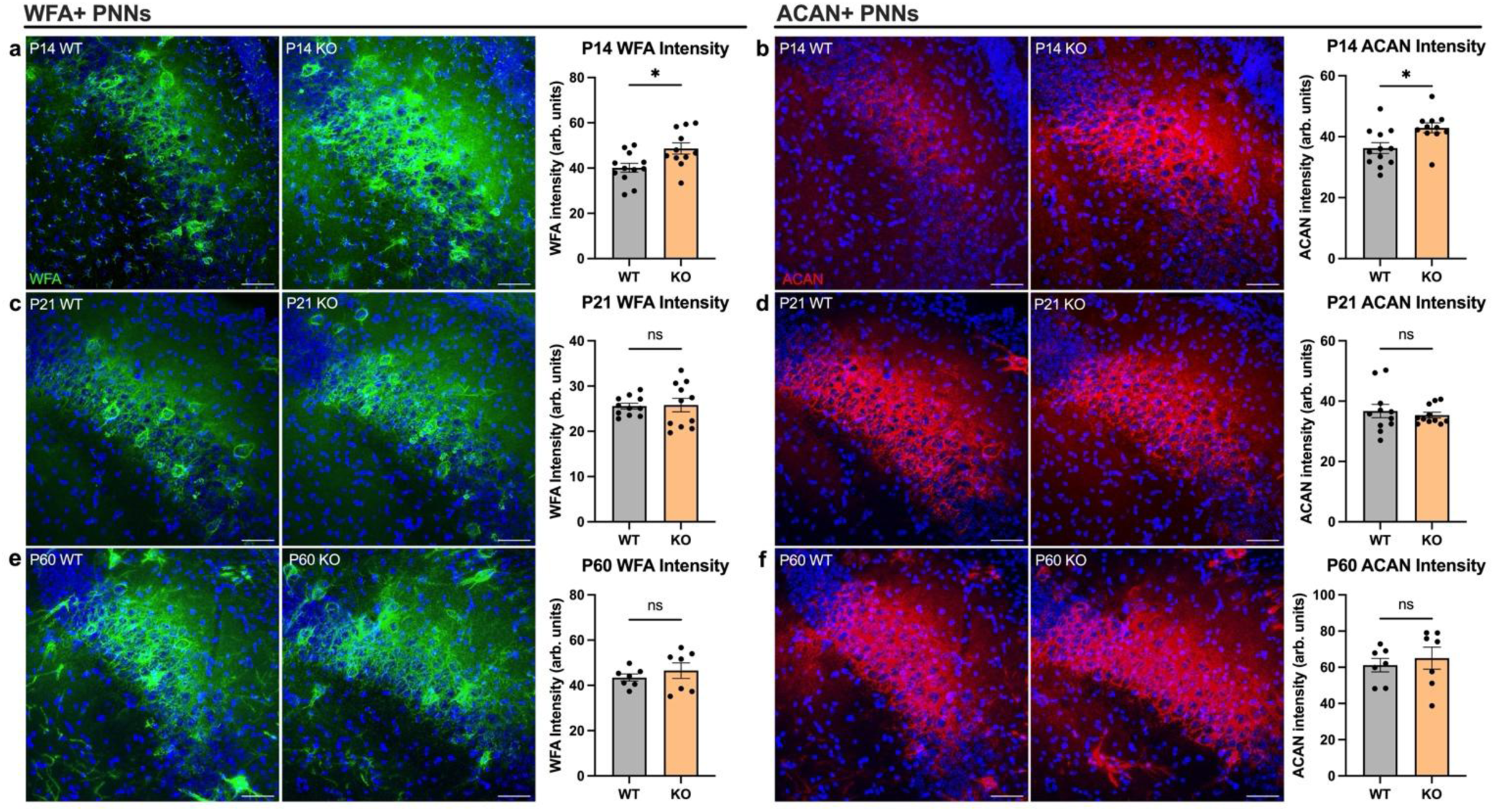
PNN intensity is increased in *Shank3B* KO mice during development. **a)** Representative CA2 images of WFA+ PNNs (green) in P14 mice of both genotypes. WFA intensity was significantly higher in P14 KO pups compared to WTs (t_21_ = 2.717, p = 0.0129; WT n = 12, KO n = 12). **b)** Representative CA2 images of ACAN+ PNNs (red) in P14 mice of both genotypes. ACAN intensity was significantly higher in P14 KO pups compared to WTs (t_21_ = 2.764, p = 0.0116). **c)** Representative CA2 images of WFA+ PNNs in P21 mice of both genotypes, which did not differ in intensity (Mann-Whitney U = 59, p = 0.9487; n = 11 per genotype). **d)** Representative CA2 images of ACAN+ PNNs in P21 mice of both genotypes, which also did not differ in intensity (Mann-Whitney U = 58, p = 0.8977). **e)** Representative CA2 images of WFA+ PNNs in P60 mice of both genotypes, which did not differ in intensity (t_12_ = 0.8142, p = 0.4314; n = 7 per genotype). **f**) Representative CA2 images of ACAN+ PNNs in P60 mice of both genotypes, which also did not differ in intensity (t_12_ = 0.5419, p = 0.5978). All images counterstained with Hoechst (blue). Scale bars = 50μm. * p<0.05; ns, not significant.

#### Reducing PNNs in the developing CA2 of *Shank3B* KO pups restores healthy social recognition into young adulthood

The degradative enzyme chondroitinase ABC (chABC) has been used to transiently degrade PNNs in the CA2 of adult mice (Hayani et al., 2018; Cope et al., 2022; Liu et al., 2023; Mehak et al., 2025). We tested different doses of chABC in the P10 CA2 that would produce a partial reduction of PNNs by P14 and, in KO pups injected with 18U/mL chABC, found that CA2 PNN intensity was significantly lower compared to those given the control enzyme, penicillinase (pnase) (t_13_ = 2.269, p = 0.0410) (Fig. 4a). After determining the appropriate chABC dose, P10 KO pups and their WT littermates received bilateral stereotaxic injections of either chABC or an identical volume of pnase directly in the CA2 region and were tested on the DSIT after a 4-day recovery period (on P14) (Fig. 4b). We found that, like unoperated WTs, those injected with pnase decreased interaction time between novel and familiar trials (p = 0.0014). Surprisingly, WT pups injected with chABC also demonstrated intact social recognition memory (p = 0.0006), possibly because our chABC treatment was designed to only partially eliminate PNNs. Like unoperated KO pups, those given pnase showed no decrease in interaction time between novel and familiar stimulus mice (p = 0.9743). In line with our hypothesis that excess PNNs contribute to impaired social recognition in KO pups, those treated with chABC with reduced PNN intensity decreased interaction time between novel and familiar exposures (p = 0.0031). Accordingly, chABC-treated KOs had a significantly higher social discrimination index than pnase-treated KOs (p = 0.0032) (Fig. 4c, d).

**Figure 4:**
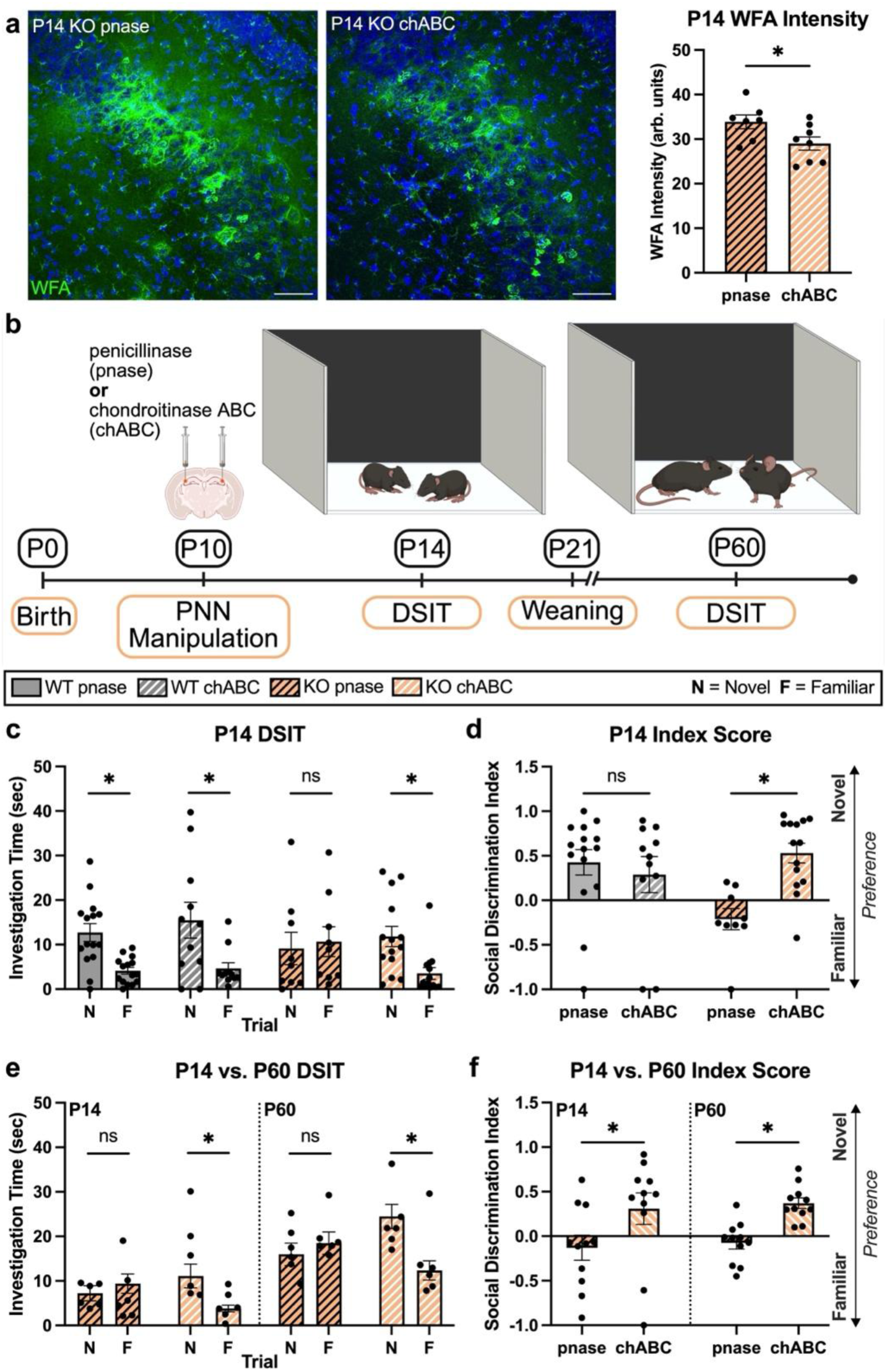
Reducing PNN expression enables social recognition in developing and adult *Shank3B* KO mice. **a)** Representative images of WFA+ PNNs (green) in the CA2 of P14 KO pups treated with the control enzyme pnase, or the degradative enzyme chABC, on P10. Images counterstained with Hoechst (blue). Scale bars = 50µm. Those treated with chABC had significantly lower WFA+ PNN intensity compared to those treated with pnase (t_13_ = 2.269, p = 0.0410; pnase n = 7, chABC n = 8). **b)** Schematic of experimental timeline. **c)** In the P14 DSIT, WT pups treated with pnase or chABC maintained preference for the novel peer over the familiar peer (pnase: p = 0.0014, n = 15; chABC: p = 0.0006, n = 11). KO pups treated with pnase remained impaired (p = 0.9743, n = 9), but those treated with chABC showed the ability to recognize and prefer the novel peer over the familiar (p = 0.0031, N = 14) (trial x treatment: F_(1, 45)_ = 5.741, p = 0.0208). **d)** KOs treated with chABC had a higher discrimination index than those treated with pnase, but no differences were seen between pnase- and chABC-treated WTs (interaction: F_(1, 45)_ = 8.547, p = 0.0054; WT p = 0.7533; KO p = 0.0032). **e)** In a separate cohort of KO mice that underwent the same manipulation, we replicated the effect on P14 where pnase mice did not discriminate between novel and familiar (p = 0.7557) while chABC mice did (p = 0.0036) (n = 11 per treatment). In adulthood, these effects persisted, with chABC mice retaining social novelty preference (p < 0.0001) and pnase mice still failing to exhibit this preference (p = 0.6409) (treatment: F_(1, 20)_ = 27.32, p < 0.0001). **f)** The P14 discrimination index was again increased in chABC mice compared to those treated with pnase (p = 0.0304), and this effect was maintained in adulthood (p = 0.0276) (treatment: F_(1, 20)_ = 11.75, p = 0.0027). * p<0.05; ns, not significant.

Next, we investigated whether the positive effect of CA2 chABC treatment on KO pups would last into adulthood or whether treated mice would eventually regress behaviorally. To address this question, a new cohort of KO mice was generated and underwent PNN manipulation on P10 in the manner described above, followed by the DSIT on P14 and again on P60. In this longitudinal experiment, we observed significant effects of treatment (chABC vs. pnase) (F_(1, 20)_ = 27.32, p < 0.0001) and of trial (novel vs. familiar) (F_(1, 20)_ = 9.985, p = 0.0049) whereby KO mice treated with chABC showed both initial and persistent improvement in social memory (Fig. 4e). On P14, pnase-treated KOs did not decrease investigation time for a novel peer (p = 0.7557). However, those treated with chABC did (p = 0.0036), replicating our findings from the previous cohort. On P60, these mice underwent the DSIT again, this time with a 24hr ITI and different novel stimulus mice. The beneficial effects of chABC persisted in KO adults, which were able to distinguish between novel and familiar social stimuli like WT mice (p < 0.0001). In contrast, KO adults treated with pnase on P10 were, as expected, persistently impaired and never gained the ability to prefer the novel peer (p = 0.6409) (Fig. 4e). In line with this finding, both P14 and P60 KO mice treated with chABC had significantly higher social discrimination indices than KO mice treated with pnase (P14: p = 0.0304; P60: p = 0.0276) (Fig. 4f). Together, the results of these experiments suggest that healthy PNN emergence in the second week of life is important for allowing social recognition behaviors to develop. KO pups exhibit unchecked accumulation of PNN components in CA2, which directly contributes to impaired social behavior that persists long after PNNs naturally lower.

#### Developmental PNN reduction increases CA2 neuronal activity and restores some aberrations in CA2 network activity in adult KO mice

Upon finding that postnatal PNN reduction improves social recognition into young adulthood in *Shank3B* KO mice, we investigated whether treatment of KO pups with chABC would reverse diminished CA2 c-fos+ cell density in response to social novelty (Fig. 2c). Indeed, chABC-treated KO mice exhibited significantly higher c-fos+ cell density in the CA2 1hr after exposure to a novel peer compared to KOs treated with pnase (t_8_ = 3.530, p = 0.0077) (Fig. 5a, b). To determine whether chABC was also sufficient to restore in vivo CA2 network activity in KOs, we implanted KO mice that were treated with chABC or pnase on P10 with a CA2 electrode in adulthood (Fig. 5c). We found that power was statistically comparable between chABC-treated and pnase-treated KO mice during bouts of immobility in theta, slow gamma, and fast gamma ranges within recording sessions (theta: F_(1, 13)_ = 0.2839, p = 0.6032; slow gamma: F_(1, 13)_ = 0.0315, p = 0.8619; fast gamma: F_(1, 14)_ = 2.026, p = 0.1765) (Fig. 5d) (Fig. S5a-c), as it was between untreated WTs and KOs in our previous experiment. Like in the baseline experiment, gamma power was transiently higher in both groups during the post-social recording session compared to recording with the novel or familiar peer (fast gamma: F_(2, 26)_ = 8.606, p = 0.0014; novel: p = 0.0453; familiar: p = 0.0011) (Fig. S5c). No differences in power were seen during active bouts of social investigation with the novel peer (F_(1, 13)_ = 1.598, p = 0.2284) (Fig. S5d). We also found that chABC treatment was ineffective at increasing slow gamma power during active investigation of a familiar partner, a metric that was diminished in untreated *Shank3B* KOs compared to untreated WTs (F_(1, 13)_ = 1.112, p = 0.3109) (Fig. S5d)

**Figure 5:**
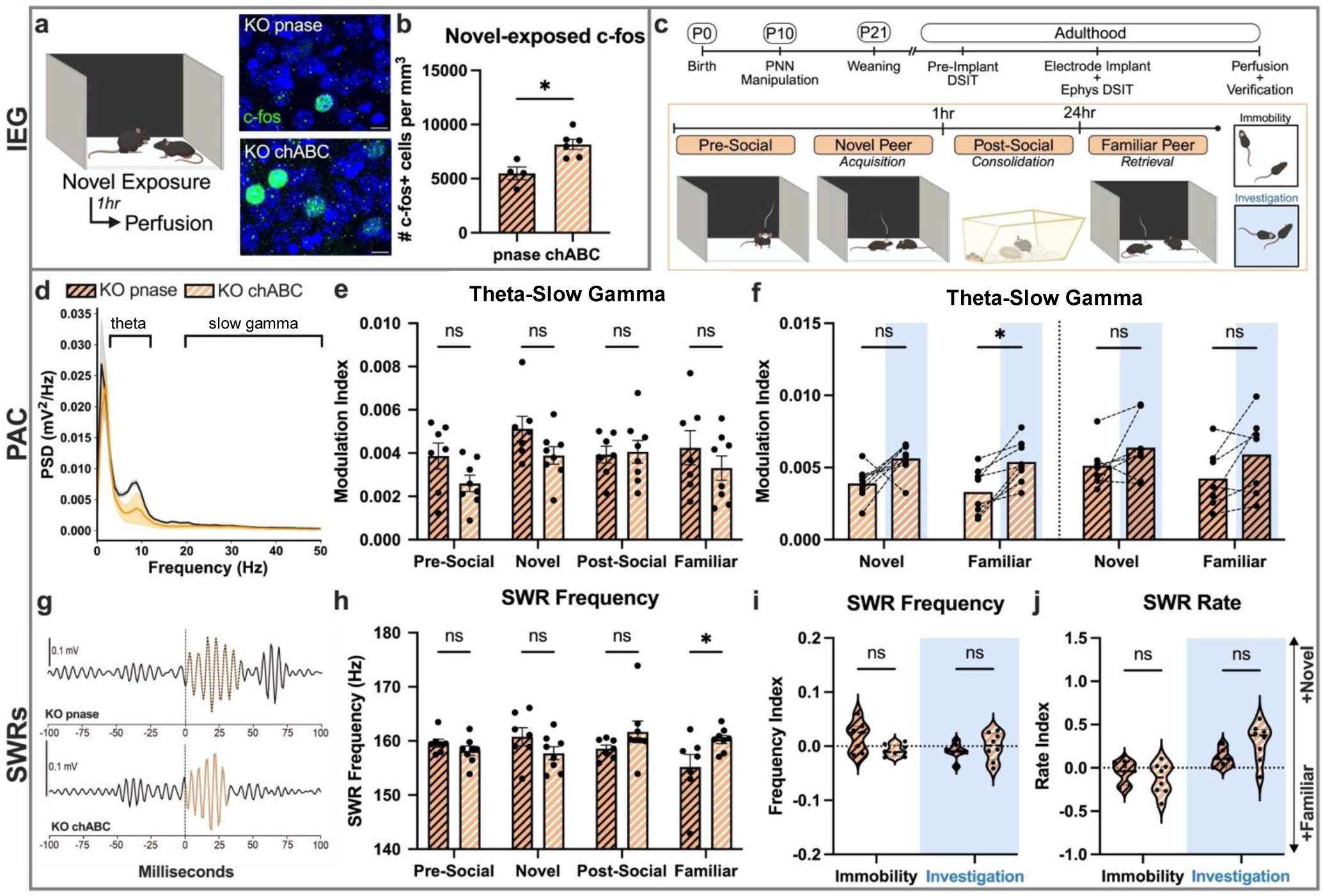
Developmental chABC treatment increases CA2 c-fos+ cell density and restores some CA2 network properties in adult *Shank3B* KO mice. **a)** Schematic of IEG experimental paradigm. **b)** KO adults treated with chABC in development had significantly higher CA2 c-fos density after exposure to a novel peer compared to those treated with pnase (t_8_ = 3.530, p = 0.0077; pnase n = 4; chABC n = 6), shown in representative images of c-fos (green) in the CA2 (counterstained with Hoechst in blue) (Scale bars = 10µm). **c)** Schematic of experimental timeline electrophysiological recording paradigm. **d)** PSD plot from representative mice highlighting theta (4-12Hz) and slow gamma (20-50Hz). **e)** PAC between theta and slow gamma was not affected based on treatment (F_(1, 13)_ = 3.516, p = 0.0834; pnase n = 7, chABC n = 8) or recording session (F_(3, 39)_ = 2.225, p = 0.1006). **f)** chABC treatment partially restored theta-slow gamma coupling. We saw a significant effect of behavioral bout type in chABC mice (F_(1, 7)_ = 51.09, p = 0.0002) where the modulation index was higher during social investigation bouts compared to immobility bouts in the familiar session (p = 0.0283), although this did not reach significance in the novel session (p = 0.0556). The same effect of bout type was seen in pnase mice (F_(1, 6)_ = 6.510, p = 0.0434) although like with untreated KOs, no post hoc differences were seen between immobility and investigation bouts in the novel (p = 0.1972) or familiar (p = 0.1018) sessions. **g)** Representative SWR trace from pnase- and chABC-treated KO mice.**h)**SWR frequency (Hz) analysis revealed a significant interaction between treatment and recording session (F_(3, 39)_ = 3.653, p = 0.0205; pre-social: p = 0.4831; post-social: p = 0.1242). Although chABC did not increase SWR frequency in the session (p = 0.1280), it was effective at increasing frequency in the familiar session (p = 0.0132). **i)**There was a significant interaction between treatment and bout type (F_(1, 13)_ = 4.741, p = 0.0485), although no post hoc effects were seen (immobility: p = 0.0549; investigation: p = 0.7059). **j)** chABC treatment did not restore the SWR rate index score during bouts of social investigation (treatment: F_(1, 13)_ = 3.440, p = 0.0030; bout type: F_(1, 13)_ = 13.26, p = 0.0865; immobility: p = 0.5946; investigation: p = 0.0775). * p<0.05; ns, not significant.

We next assessed theta-slow gamma PAC and, like in the previous experiment, found no differences in the modulation index within recording sessions during bouts of immobility (Fig. 5e). However, PAC during active investigation appeared to be partially restored in chABC-treated KOs and, like untreated WTs, mice in this group exhibited a control-like increase in PAC during active investigation of a social partner compared to during bouts of immobility within the same session (F_(1, 7)_ = 51.09, p = 0.0002) (Fig. 5f). However, treatment was only effective at restoring this increase during investigation of the familiar peer (p = 0.0283) but was not sufficient to produce a statistically significant increase during investigation of the novel peer (p = 0.0556). KO mice given pnase demonstrated a significant effect of bout type on PAC but, like untreated KOs, failed to modulate coupling as a factor of investigation in either session (F_(1, 6)_ = 6.510, p = 0.0434; novel: p = 0.1972; familiar: p = 0.1018) (Fig. 5f).

Lastly, we analyzed SWR properties that were atypical in untreated KO mice to determine whether they were changed in adults that underwent P10 chABC treatment (Fig. 5g). SWR frequency, which was low in unoperated KOs during both the novel and familiar exposures, did not differ between chABC-treated and pnase-treated KOs during exposure to the novel peer (p = 0.1280) (Fig. 5h). The following day, however, chABC-treated mice exhibited a significant increase in SWR frequency during exposure to the familiar peer compared to pnase-treated mice, indicating that treatment successfully prevented decreased SWR frequency seen in untreated KOs (p = 0.0132) (interaction: F_(3, 39)_ = 3.653, p = 0.0205 (Fig. 5h). Like in untreated KOs and WTs, pre- and post-social SWR frequency did not differ between groups (pre-social: p = 0.4831; post-social: p = 0.1242) (Fig. 5h). Unlike in the baseline experiment, here we observed a significant interaction between treatment and session in the SWR frequency index score, although no post hoc differences were detected (interaction: F_(1, 13)_ = 4.741, p = 0.0485; immobility: p = 0.0549; investigation: p = 0.7059) (Fig. 5i). As an index score, SWR rate was previously diminished in untreated KOs only during active social investigation and not during bouts of immobility, suggesting fewer SWRs than WTs during exposure to a novel partner (Fig. 2l). Treatment did not significantly increase the rate index during investigation in chABC-treated KOs (p = 0.0775); however, these mice on average exhibited a twofold higher index than pnase-treated controls during active investigation, suggesting more SWRs during novel social exposure (treatment: F_(1, 13)_ = 3.440, p = 0.0030; bout type: F_(1, 13)_ = 13.26, p = 0.0865; immobility: p = 0.5946) (Fig. 5j). Our results suggest that developmental PNN reduction in *Shank3B* KO mice may increase CA2 neuronal activity during exposure to a novel peer, and prevent some atypical CA2 oscillatory activity, including in PAC and SWRs during exposure to a familiar peer, and these changes may undergird improvements in social recognition behavior seen after chABC treatment.

### PNN-associated semaphorin-3A and afferent input to CA2 are atypical in Shank3B KO mice

Since excess CA2 PNN accumulation in the early postnatal period negatively affects hippocampus-dependent social recognition and network activity in *Shank3B* KO mice, we next investigated a potential mechanism by which aberrant PNNs might induce dysfunction. Semaphorin-3A (sema3A) is often associated with PNNs and mediates axon ingrowth and circuit formation in the developing brain (Uesaka et al., 2014; Fawcett et al., 2019), leading us to investigate whether these processes were atypical in the CA2 of *Shank3B* KO mice in the second postnatal week, when preference for social novelty fails to emerge.

#### Developing *Shank3B* KO mice exhibit excess PNN-associated semaphorin-3A, which is reduced after chABC treatment

We labeled P14 WT and KO tissue with both sema3A and WFA and confirmed that both genotypes exhibited a positive relationship between intensity of these markers (r = 0.6890, p = 0.0275) (Fig. 6a, b). Area CA2 in P14 KO mice contained significantly higher intensity and distribution of sema3A labeling compared to WTs (intensity: t_8_ = 3.436, p = 0.0089; percent area: t_8_ = 3.254, p = 0.0116) (Fig. 6c, d). In P14 pups that underwent P10 PNN manipulation, a positive relationship was still apparent in sema3A and WFA labeling (r = 0.5196, p = 0.0001) but the intensity and distribution of sema3A staining was significantly diminished in KO pups treated with chABC compared to those given pnase, suggesting that reducing CA2 PNNs during development prevents excess accumulation of sema3A (intensity: t_21_ = 2.527, p = 0.0196; percent area: t_21_ = 2.347, p = 0.0288) (Fig. 6b-d). Given these findings, we next assessed whether projections to the developing CA2 might also be atypical in young and adult *Shank3B* KO mice.

**Figure 6:**
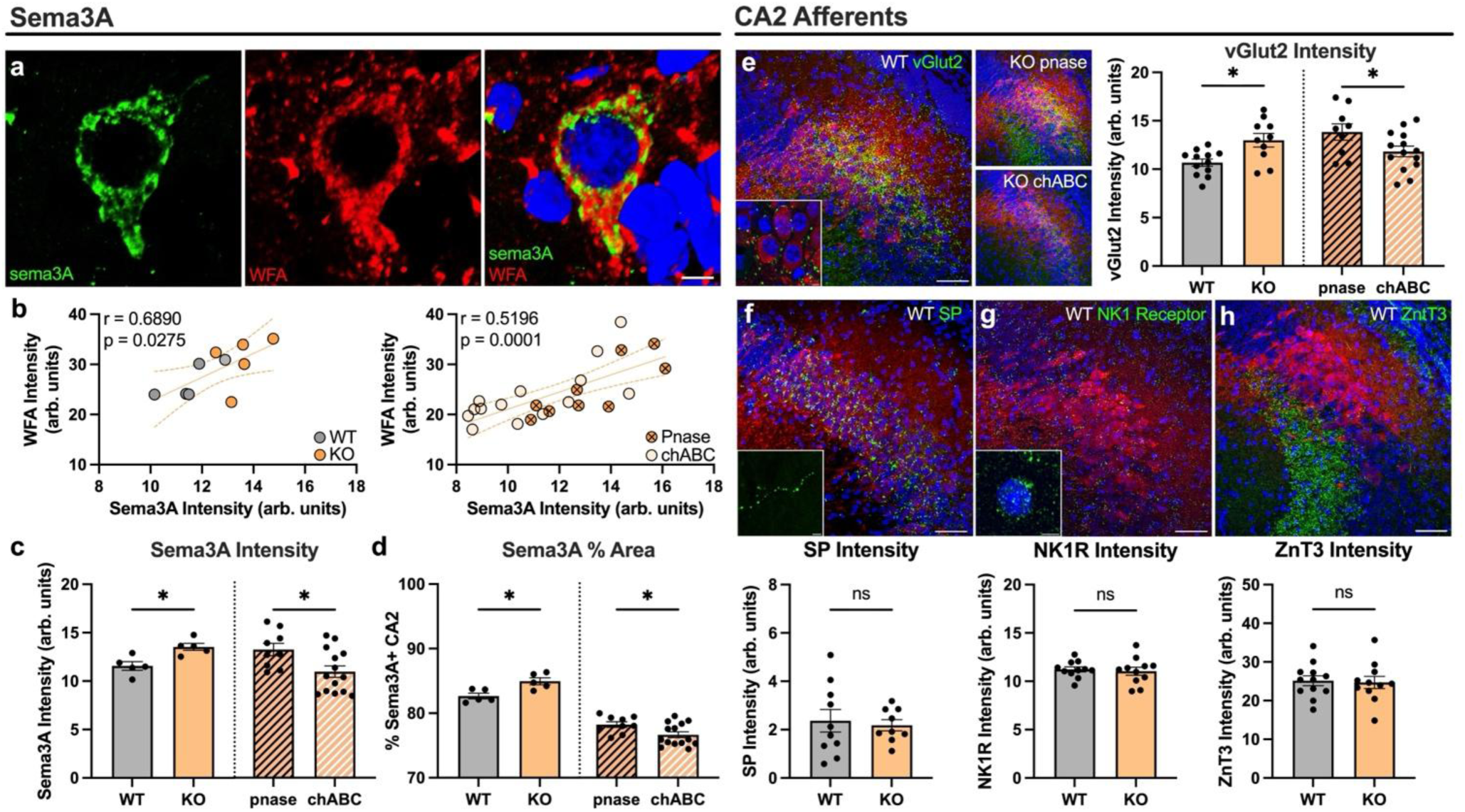
Semaphorin-3A is increased in P14 *Shank3B* KO mice, potentially altering glutamatergic SuM-CA2 signaling. **a)** Representative P14 image of a sema3A+ (green) and WFA+ (red) CA2 neuron in a P14 KO mouse, counterstained with Hoechst (blue). Scale bar = 10µm. **b)** Sema3A+ labeling intensity was positively correlated with WFA intensity in both unmanipulated (r = 0.6890, p = 0.0275; n = 5 per genotype) and P10 operated mice (r = 0.5196, p = 0.0001; pnase n = 9, chABC n = 14) (regression line shown). **c)** Sema3A intensity was increased in P14 KOs (t_8_ = 3.436, p = 0.0089), and this was normalized in mice that underwent P10 chABC treatment (t_21_ = 2.527, p = 0.0196). **d)** The percent area of CA2 labeled with sema3A was significantly increased in baseline P14 KO mice compared to WTs (t_8_ = 3.254, p = 0.0116), and this was lowered with chABC treatment (t_21_ = 2.347, p = 0.0288). **e)** Representative images of vGlut2 (green) in the CA2 (red) of a P14 WT pup as well as pnase- and chABC-treated KO mice (scale bar = 50μm and applies to all low magnification images). P14 vGlut2 intensity was increased in KOs compared to WTs (t_20_ = 3.062, p = 0.0062; WT n = 12, KO n = 10), and this was restored in chABC-treated KOs (t_21_ = 2.094, p = 0.0485). **f)** Representative images from a WT mouse of SP labeling (green) in the CA2 (red). SP labeling intensity was comparable between genotypes at P14 (Mann-Whitney U = 44, p = 0.9682; WT n = 10, KO n = 9). **g)** Representative P14 image of NK1R labeling (green) in the CA2 (red). Labeling intensity of the NK1R, which is the active receptor of SP, was typical at P14 (t_20_ = 0.3732, p = 0.7129; n = 11 per genotype). **h)** Representative image of ZnT3 labeling (green) in the P14 WT CA2 (red). ZnT3+ innervation from the DG was comparable between P14 KOs and WTs (t_21_ = 0.2139, p = 0.8327; WT n = 12, KO n = 11). All images counterstained with Hoechst (blue) and CA2 identified with RGS14 (red). Scale bars = 50μm (low magnification) or 10μm (high magnification). * p<0.05; ns, not significant.

#### SuM afferents to the CA2 appear to be overgrown in developing *Shank3B* KO mice, and are reduced by chABC treatment

Next, we labeled axons from the supramammillary nucleus of the hypothalamus (SuM), which are the source of vGLUT2 in the CA2 (Halasy et al., 2004), as well as the mossy fibers from dentate gyrus, which are the source of ZnT3 in the CA2 (Wenzel et al., 1997), since both projections are present by P14 and participate in social recognition memory in adult mice (Chen et al., 2020; Laham et al., 2022; Diethorn and Gould, 2023a). We found that the intensity of vGlut2 was significantly higher in P14 KO mice compared to WT controls (t_20_ = 3.062, p = 0.0062) (Fig. 6e), but no differences between genotypes were seen in ZnT3 intensity (t_21_ = 0.2139, p = 0.8327) (Fig. 6h). Since a subset of SuM axons release the neuropeptide substance P (SP) which binds to neurokinin 1 receptors (NK1Rs) on CA2 neurons (Dasgupta et al., 2017), we also examined the intensity of SP and NK1R labeling and found no differences between *Shank3B* KOs and WTs at P14 (SP: Mann-Whitney U = 44, p = 0.9682; NK1R: t_20_ = 0.3732, p = 0.7129) (Fig. 6f, g). Because sema3A is known to inhibit synapse elimination (Uesaka et al., 2014), we hypothesized that high levels of PNNs with their excess sequestered sema3A may be preventing the elimination of vGlut2+ SuM axons. To determine whether excessive PNNs are responsible for high vGlut2 intensity, we examined this measure in *Shank3B* KO pups treated with chABC on P10 and found a reduction in vGlut2 intensity by P14 compared to KO pups treated with pnase (t_21_ = 2.094, p = 0.0485) (Fig. 6e).

#### Some measures of SuM-CA2 circuitry normalize in *Shank3B* KOs by adulthood, but new aberrations arise

Although untreated adult *Shank3B* KO mice exhibit persistent impairments in social memory during young adulthood, we found that the high intensity of vGlut2-expressing SuM input seen at P14 was no longer present in KO mice at P60 (t_18_ = 0.5693, p = 0.5762) (Fig. S6a). In contrast to results found on P14, however, innervation from SP-expressing SuM axons appeared to be aberrant in adult *Shank3B* KO mice. In the CA2, the average size of SP+ boutons was significantly larger in KO mice compared to WT controls, although overall labeling intensity was not affected (size: t_18_ = 2.817, p = 0.0114; WT: 0.72μm^2^ ± 0.0352; KO: 0.88μm^2^ ± 0.0445) (intensity: t_18_ = 1.249, p = 0.2277) (Fig. S6b, c). In addition, labeling intensity of the NK1 receptor, which is the active receptor of SP, was significantly diminished in the CA2 of adult KOs (t_18_ = 2.682, p = 0.0152) (Fig. S6d). These findings suggest that despite normalization of some measures of SuM afferents between P14 and adulthood, the *Shank3B* KO CA2 exhibits aberrations that dynamically vary across development and may contribute to persistent social recognition impairments.

As a final probe of the SuM in *Shank3B* KO mice, we assessed the responsiveness of SuM to novel social stimuli using c-fos, since others have demonstrated that the SuM-CA2 circuit responds preferentially to social novelty (Chen et al., 2020). We first confirmed that c-fos+ cell density in SuM was not different between adult KOs and WTs perfused directly from the home cage (p = 0.9592) (Fig. S6e-g). In a separate cohort of adult WTs and KOs perfused 1hr after exposure to a novel peer, SuM neuronal activation was modulated by social novelty, and c-fos+ cell density was significantly elevated (WT: p < 0.0001; KO: p < 0.0001). Novel-exposed KO mice, however, had significantly lower c-fos+ cell density in SuM compared to novel-exposed WTs, paralleling our findings from CA2 (interaction: F_(1, 29)_ = 9.619, p = 0.0043; novel-exposed: p = 0.0007) (Fig. S6f). Because the SuM has reciprocal connections with the CA2 (Cui et al., 2013), the SuM c-fos effect may be secondary, resulting from atypical projections of the CA2. Taken together, these findings suggest that PNNs and associated sema3A abnormalities in the developing CA2 disorganize incoming SuM axons, and that continued sculpting of this circuit contributes to social impairments in adulthood despite some circuit-level restoration.

## Discussion

The results of these experiments suggest that social recognition deficits observed in developing and adult *Shank3B* KO mice arise from excessive PNNs in the CA2 region, which sequester higher quantities of sema3A, potentially leading to delayed pruning of SuM inputs to the CA2. Reduction of CA2 PNNs during the postnatal period restores social memory function and some aberrant network activity in *Shank3B* KO adults, potentially by reducing sema3A and restoring SuM inputs to WT levels during a critical period in development. These findings suggest that specific behavioral deficits of a genetic mutation associated with neurodevelopmental disorders can be lastingly corrected with early life intervention.

We found that many behaviors were typical in *Shank3B* KO mice at several developmental timepoints, including isolation-induced USVs on P5, long-term social recognition on P14, and object location memory on P21 and P60, but KO mice exhibited specific deficits in social novelty preference and short-term social recognition during development and in adulthood when most healthy mice demonstrate this ability. These results strongly suggest that mutation or loss of Shank3, a synaptic scaffolding protein, does not disrupt development of circuits associated with social or hippocampal function in general, but rather produces a specific deficit that impairs the CA2 region. In healthy mice, hippocampal area CA2 is necessary for the ability to recognize and prefer novel peers both during development and in adulthood (Hitti and Siegelbaum, 2014; Stevenson and Caldwell, 2014; Diethorn and Gould, 2023a). This ability is in part supported by the presence of PNNs, which begin to emerge in CA2 during the second postnatal week (Carstens et al., 2016; Diethorn and Gould, 2023a,b). Atypical PNNs in CA2 have been characterized in adult mouse models for neurodevelopmental conditions and Alzheimer’s disease that exhibit social memory dysfunction (Carstens et al., 2021; Cope et al., 2022; Rey et al., 2022). We found that CA2 PNNs were atypical in developing *Shank3B* KO mice, which expressed excess PNN labeling compared to healthy WT controls on P14. This effect was not observed at weaning nor in young adulthood, but nevertheless social dysfunction persisted in adult *Shank3B* KO mice and was accompanied by impairments in CA2 neuronal activation, as well as in PAC and SWRs. In adult mice that underwent developmental PNN reduction, social dysfunction was prevented and CA2 neuronal activation, as well as PAC and SWRs, during social experience were partially improved. These findings suggest that area CA2 may undergo a sensitive period of plasticity mediated by PNN emergence during postnatal development and that uncorrected PNN abnormalities during this time can persistently disorganize the region.

Healthy social recognition abilities also require the ventral CA1, a major target site of CA2 neurons (Okuyama et al., 2016). Studies have shown that adult *Shank3B* KO mice exhibit diminished activation of vCA1 neuronal ensembles in response to social experience (Tao et al., 2022), and that chemogenetic activation of the CA2-vCA1 pathway can temporarily correct social recognition abilities (Cope et al., 2023). In *Shank3* mutants, diminished neuronal activity in vCA1 may be secondary to more direct effects on the CA2, but conditional knockout of *Shank3* in vCA1 neurons alone can also impair social memory (Chung et al., 2024). Taken together, these findings suggest that transgenic *Shank3* mutation directly impacts both CA2 and vCA1, either of which may be sufficient to produce social dysfunction. Despite these abnormalities, social recognition function can be restored temporarily by activating the CA2 -vCA1 circuit in adulthood or lastingly by lowering CA2 PNNs during development. The extent to which developmental CA2 PNN correction affects vCA1 function remains unexplored.

We considered the relationship between PNNs and the axon guidance cue molecule semaphorin-3A, since it is sequestered within PNNs and influences local microcircuitry by mediating synapse growth and elimination (Vo et al., 2013; Uesaka et al., 2014; Boggio et al., 2019), and found that excess PNNs in *Shank3B* KO pups were accompanied by increased sequestration of sema3A, which may impede developing circuitry. Indeed, vGlut2+ innervation from the supramammillary nucleus (SuM) was also increased in P14 *Shank3B* KO pups. This abnormality may contribute to the observed behavioral impairments due to the role of this pathway in social recognition in adulthood (Chen et al., 2020). Infusing chABC into the CA2 on P10 lowered PNN levels in KO pups by P14, which was accompanied by lowered sema3A and vGlut2 expression. It appears, however, that excess sema3A expression in P14 KOs did not disrupt zinc transporter 3+ mossy fiber input from the dentate gyrus to CA2. It may be the case, then, that the role of PNN-associated sema3A in guiding afferent input does not broadly affect all afferent populations and only specifically impacts some developing innervation, namely from the SuM. Although measures of vGLUT2 innervation normalized by adulthood in unoperated *Shank3B* KOs, other measures that reflect a subset of this projection known to release SP, appear abnormal, changes that seem to occur late perhaps due to compensatory mechanisms. These differences, along with diminished activation of SuM neurons in response to social novelty in *Shank3B* KOs, suggest persistent abnormalities in the SuM-CA2 circuit in adulthood. Since SuM inputs to the hippocampus have been linked to neuronal oscillations, including theta rhythm (Vertes et al., 2015) and SWRs (Vicente et al., 2020), persistently diminished activity in this afferent population may play a substantial role in electrophysiological and behavioral dysfunction in *Shank3B* KO adults,

Collectively, these findings demonstrate that excess PNNs and associated molecular changes in CA2 disrupt the maturation of social recognition circuits in *Shank3B* KO mice. By identifying a postnatal window during which PNN emergence shapes CA2 connectivity and behavior through sema3A signaling, our work highlights a developmental mechanism that may underlie both early and persistent social impairments. Future studies aimed at dissecting the developmental timing of SuM and other afferent pathways to the CA2 will be essential for understanding how early-life circuit miswiring leads to long-term social dysfunction and may be informative about interventions for neurodevelopmental disorders.

## Acknowledgments

The authors thank Drs.Timothy Buschman and Polina Iamshchinina for helpful advice about electrophysiological analyses. This work was supported by the National Institutes of Health, NIMH R01 MH118631 (EG) and the Princeton University Charlotte Elizabeth Proctor Honorific Fellowship (EJD).

---

**Figure S1:**
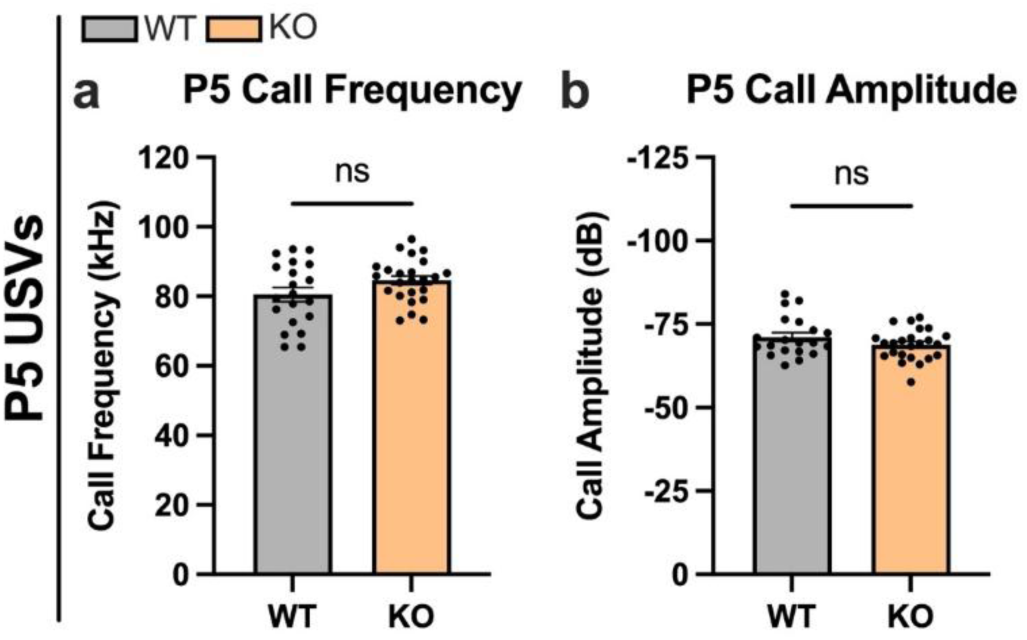
USV amplitude and frequency are similar between genotypes on P5. **a)** No effect of genotype was seen in average frequency of USVs (kHz) emitted in isolation (t_42_ = 1.737, p = 0.0897; WT n = 20, KO n = 24). **b)** No effect of genotype was seen in USV call amplitude (dB) in isolation (t_42_ = 1.431, p = 0.1597). ns, not significant.

**Figure S2:**
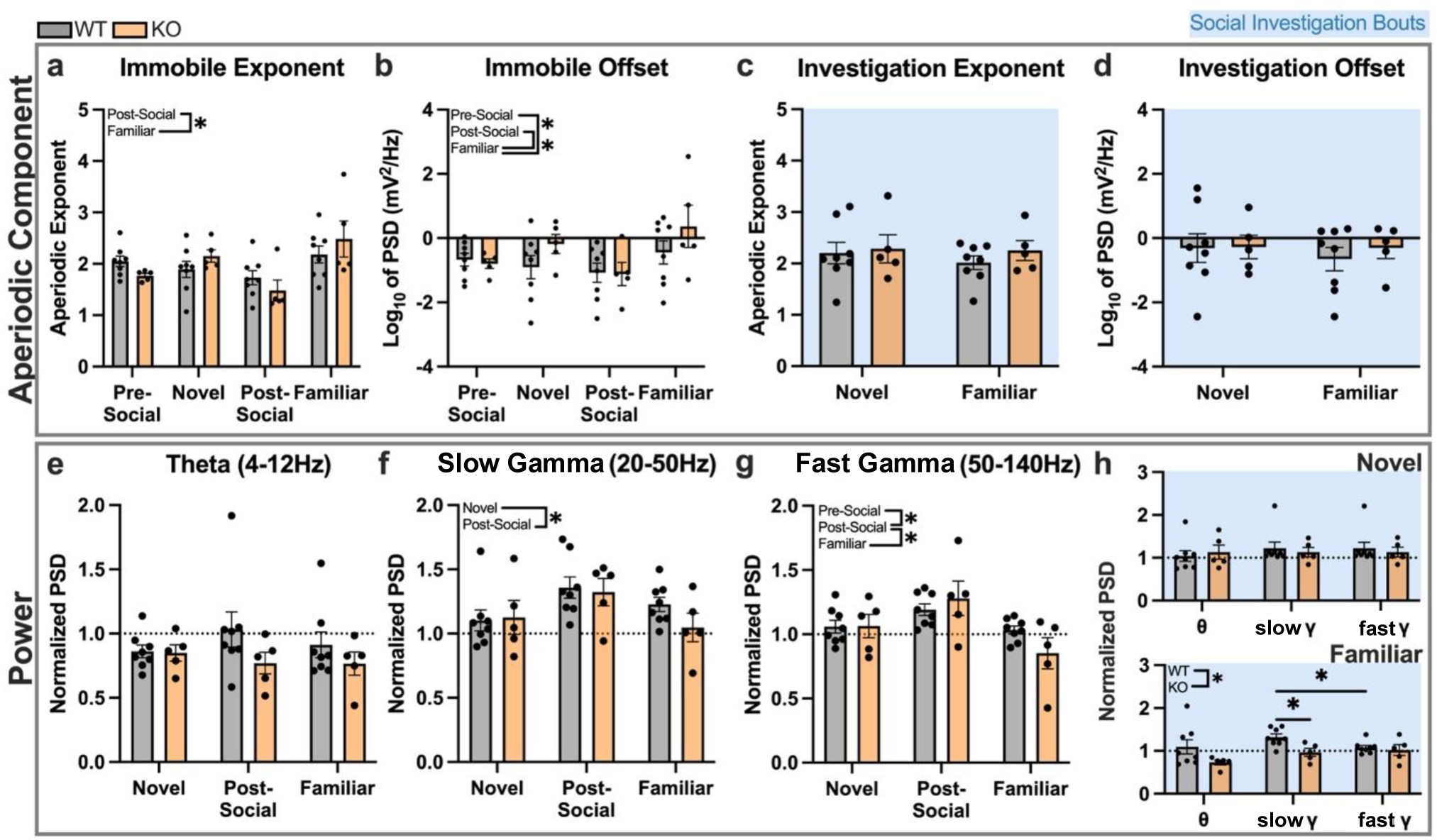
*Shank3B* KO adults do not exhibit differences in the CA2 aperiodic component, but some PSD measures slightly deviate from WTs. **a)** During periods of immobility away from the stimulus mouse, both WTs and KOs had a higher aperiodic exponent (slope) with the familiar peer compared to the post-social recording (session: F_(2.299, 25.29)_ = 7.000, p = 0.0027, E = 0.7663; post-social vs. familiar: p = 0.0267; WT n = 8, KO n = 5). **b)** The same effect was seen with the aperiodic offset (intercept) (session: F_(3, 33)_ = 6.796, p = 0.0011; post-social vs. familiar: p = 0.0006), and this was also true when comparing the pre-social and familiar sessions (p = 0.0391). **c)** No differences were found during active bouts of social investigation in the aperiodic exponent (trial: F_(1, 11)_ = 0.7153, p = 0.4157; genotype: F_(1, 11)_ = 0.3810, p = 0.5496) nor in the **d)** offset (session: F_(1, 11)_ = 0.8680, p = 0.3715; genotype: F_(1, 11)_ = 0.1199, p = 0.7356). **e)** No effects were found during periods of immobility in normalized theta power (trial: F_(1.595, 17.54)_ = 0.2023, p = 0.7697, E = 0.7975; genotype: F_(1, 11)_ = 2.675, p = 0.1302). **f)** A significant effect of session (F_(1.928, 21.21)_ = 6.250, p = 0.0078, E = 0.9640), but not of genotype (F_(1, 11)_ = 0.3833, p = 0.5484), was seen where slow gamma power in the post-social recording was significantly higher than that with the novel peer (p = 0.0077). **g)** An interaction between trial and genotype was observed in fast gamma power, and PSD during post-social consolidation was higher than during interaction with the novel or familiar mouse (F_(1.565, 17.21)_ = 5.069, p = 0.0248, E = 0.7823; novel: p = 0.0035; familiar: p = 0.0038). WTs also decreased power with the familiar peer compared to the post-social recording conducted the previous day (p = 0.0443), a comparison that did not reach significance in KOs (p = 0.0547). **h)** During bouts of social investigation, no differences were seen in power between genotypes in the novel trial in theta, slow gamma, nor fast gamma ranges (band: F_(2, 22)_ = 1.212, p = 0.3168; genotype: F_(1, 11)_ = 0.0227, p = 0.8830). With the familiar peer there was a significant effect of genotype, and slow gamma power, but not theta nor fast gamma power, was significantly diminished in KOs compared to WTs (genotype: F_(1, 11)_ = 6.085, p = 0.0313; theta: p = 0.0683; slow gamma: p = 0.0154; fast gamma: p = 0.6907). Slow gamma power also dominated over fast gamma power in the WTs (p = 0.0203), but not in KOs (p = 0.8347), and no effect of band was seen (F_(1.255, 13.81)_ = 2.572, p = 0.1268, E = 0.6227). * p<0.05.

**Figure S3:**
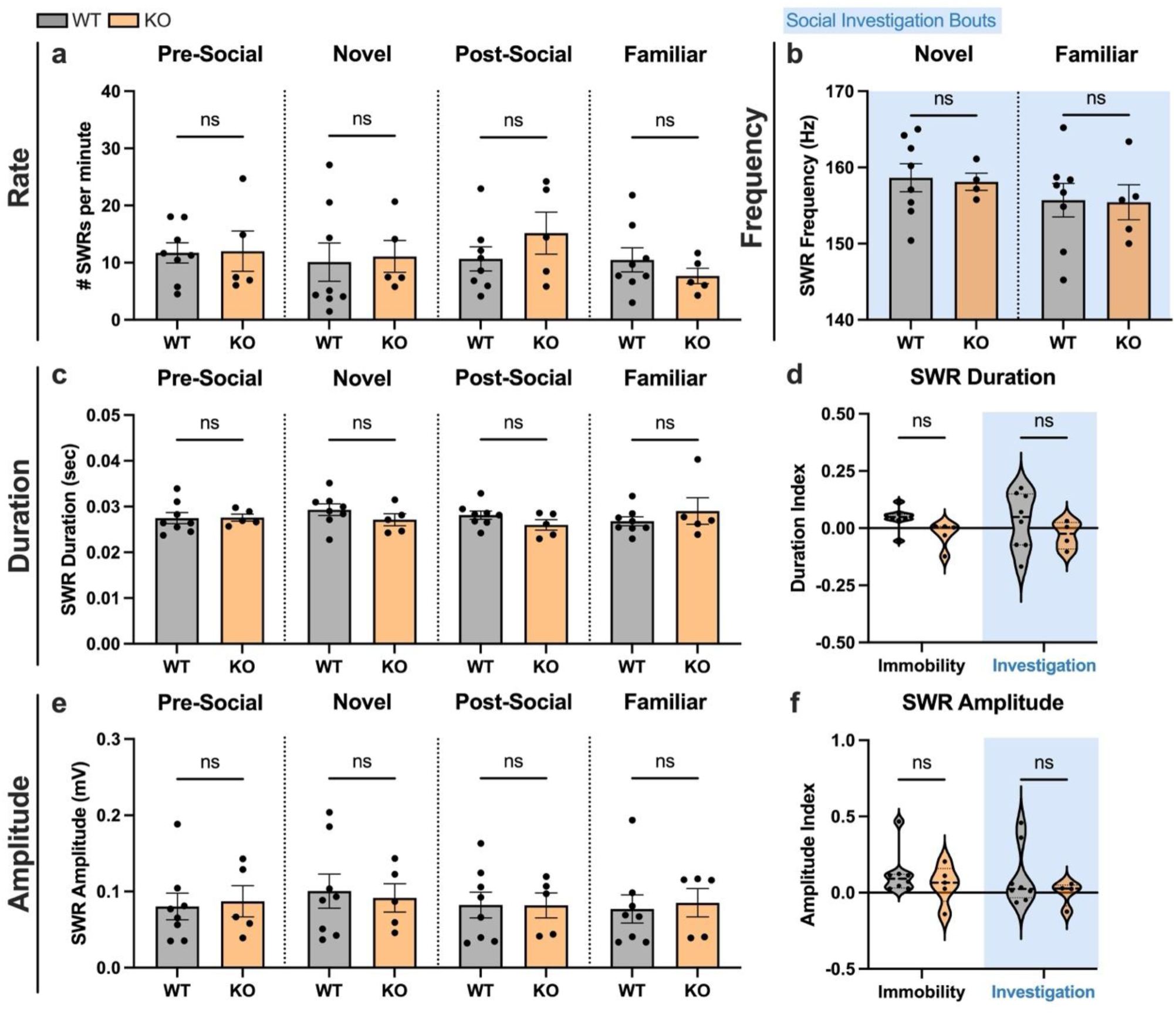
Sharp wave ripple properties are similar between *Shank3B* KO and WT adults. **a)** No differences were found in SWR rate (# SWRs/min. immobile) within recording sessions (pre-social: t_11_ = 0.0844, p = 0.9342; novel: t_11_ = 0.2076, p = 0.8393; post-social: t_11_ = 1.147, p = 0.2756; familiar: t_11_ = 0.9603, p = 0.3575; WT n = 8, KO n = 5). **b)** SWR frequency (Hz) was not increased during active social investigation with either stimulus (novel: t_10_ = 0.1923, p = 0.8514; familiar: t_11_ = 0.0808, p = 0.9370). **c)** SWR duration (sec) was consistent between genotypes in each recording session (pre-social: t_11_ = 0.0631, p = 0.9508; novel: t_11_ = 1.146, p = 0.2760; post-social: t_11_ = 1.472, p = 0.1692; familiar: t_11_ = 0.8728, p = 0.4014). **d)** No effects were seen in the duration index score based on genotype or behavioral bout (interaction: F_(1, 21)_ = 0.0169, p = 0.8975). **e)** Like with rate and duration, no differences were found in SWR amplitude (mV) within recording sessions (pre-social: t_11_ = 0.2460, p = 0.8102; novel: t_11_ = 0.2762, p = 0.7875; post-social: t_11_ = 0.0217, p = 0.9831; familiar: t_11_ = 0.2931, p = 0.7749). **f)** No effects were seen in the SWR amplitude index score (interaction: F_(1, 10)_ = 0.0143, p = 0.9072). ns, not significant.

**Figure S4:**
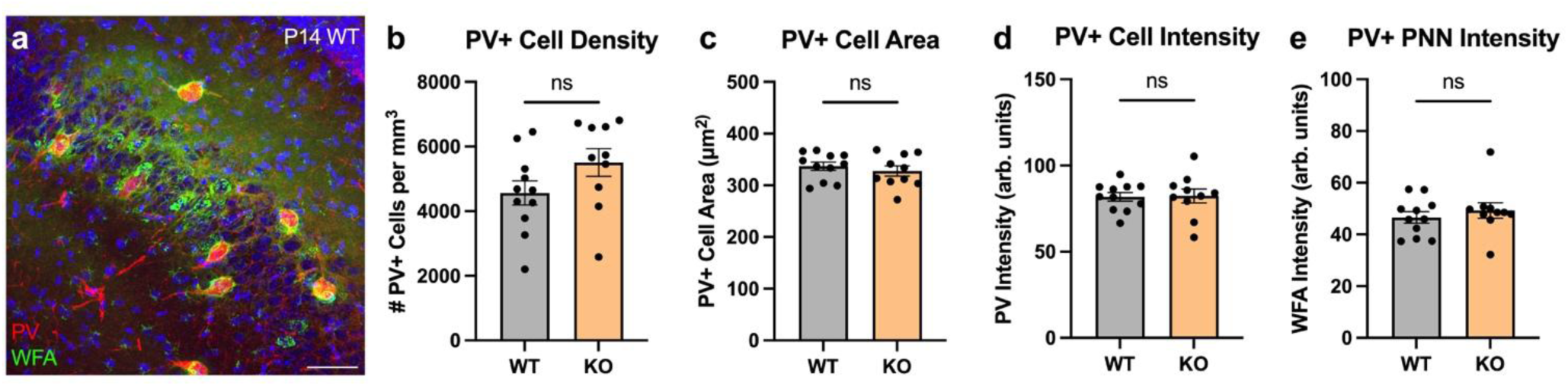
PV+ interneuron measures are similar between P14 *Shank3B* KO and WT mice. **a)** Representative image from a P14 WT of PV (red) and WFA (green) in the CA2, counterstained with Hoechst (blue). Scale bar = 50μm. **b)** The density of PV+ cells in CA2 was not atypical in P14 KO mice (t_19_ = 1.668, p = 0.1117; WT n = 11, KO n = 10). **c)** The average area of PV+ cells was also similar between genotypes (t_19_ = 0.7301, p = 0.4742). **d)** The intensity of PV+ cells in CA2 was comparable between KO and WT mice at P14 (t_19_ = 0.1213, p = 0.9047). **e)** The intensity of WFA+ PNNs surrounding PV+ interneurons was not affected in P14 KO mice (t_19_ = 0.7416, p = 0.4674). ns, not significant.

**Figure S5:**
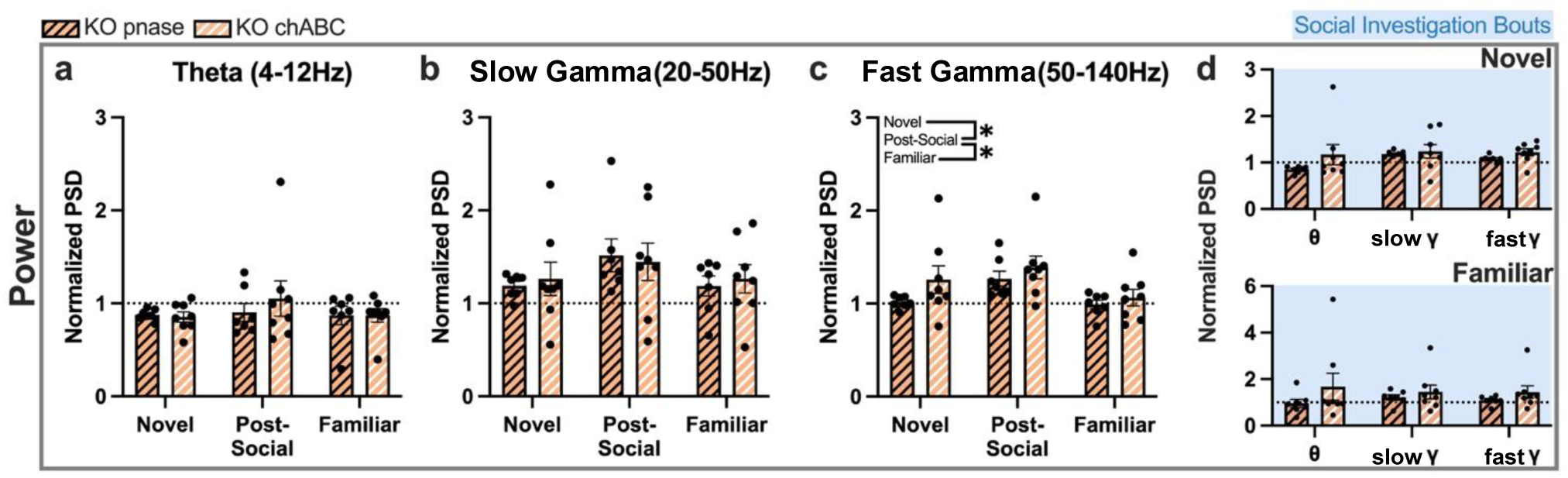
chABC-treated KO adults exhibit similar CA2 power compared to those treated with the control enzyme. **a)** No effects of recording session (F_(2, 26)_ = 0.6342, p = 0.5384) or of treatment (F_(1, 13)_ = 0.2839, p = 0.6032) were seen in normalized theta power during bouts of immobility, which was also the case in **b)** slow gamma power (session: F_(2, 26)_ = 2.223, p = 0.1284; treatment: F_(1, 13)_ = 0.0315, p = 0.8619). **c)** In the fast gamma range, a significant effect of recording session was observed (F_(2, 26)_ = 8.606, p = 0.0014) whereby power was higher in the post-social consolidation period than with the novel peer (p = 0.0011) or with the familiar peer (p = 0.0435). **d)** No effects of treatment were seen in normalized power during active social investigation during either exposure (novel: F_(1, 13)_ = 1.598, p = 0.2284; familiar: F_(1, 13)_ = 1.112, p = 0.3109). *p<0.05.

**Figure S6:**
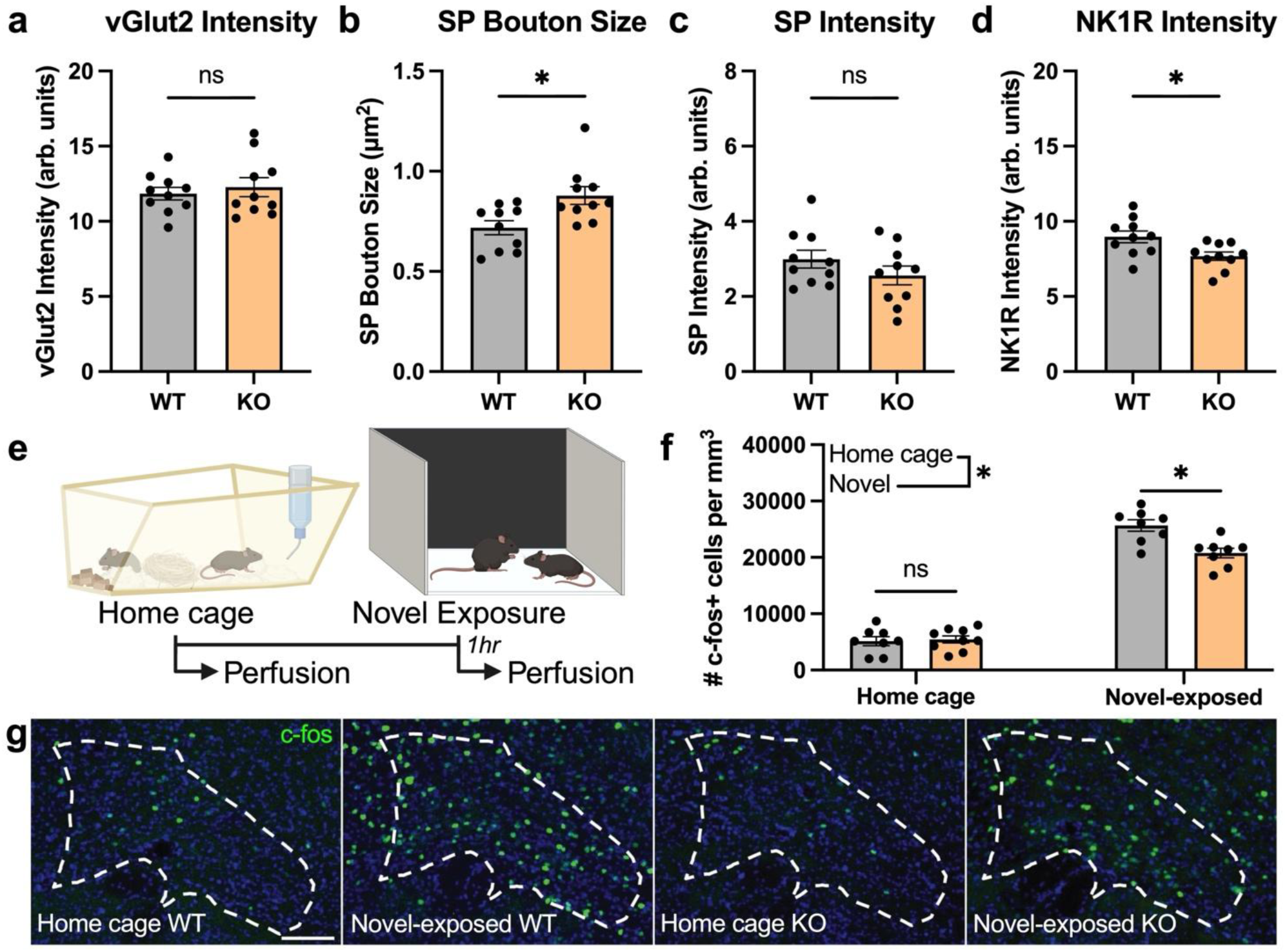
SuM and SuM-CA2 aberrations are apparent in adult *Shank3B* KO mice. **a)** Although increased in P14 KOs, CA2 vGlut2 intensity was comparable between P60 WTs and KOs (t_18_ = 0.5693, p = 0.5762; n = 10 per genotype). **b)** The average size of SP+ boutons was increased in P60 KO mice compared to controls (t_18_ = 2.817, p = 0.0114; n = 10 per genotype), although the intensity of SP **(c)** was unaffected (t_18_ = 1.249, p = 0.2277). **d)** Within the same KO tissue, there was a corresponding decrease in labeling intensity for the NK1R, which is the active receptor of SP, compared to WT controls (t_18_ = 2.682, p = 0.0152). **e)** Schematic of IEG experimental paradigm. **f)** In mice perfused from the home cage, no differences were seen in SuM c-fos+ cell density between KOs and WTs (p = 0.9592; WT n = 9, KO n = 10). In mice perfused after exposure to a novel peer, cell density was significantly increased compared to home cage counterparts (WT: p < 0.0001, n = 9; KO: p < 0.0001, n = 9), but novel-exposed KOs exhibited significantly fewer c-fos+ cells compared to novel-exposed WTs (interaction: F_(1, 29)_ = 9.619, p = 0.0043; p = 0.0007). **g)** Representative images of c-fos+ cells (green) in the lateral SuM in adult WT and KO mice perfused from the home cage and 1hr after exposure to a novel peer. Images counterstained with Hoechst (blue). Scale bars = 100μm. *p<0.05; ns, not significant.

## Notes

### Competing Interest Statement

The authors have declared no competing interest.

### Summary of Updates

The new version has a modified title, text edits to the electrophysiology sections, and discussion added.

## References

Alexander, G. M., Nikolova, V. D., Stöber, T. M., Gruzdev, A., Moy, S. S., & Dudek, S. M. (2025) Perineuronal nets on CA2 pyramidal cells and parvalbumin-expressing cells differentially regulate hippocampal-dependent memory. Journal of Neuroscience, 45(6). 10.1523/JNEUROSCI.1626-24.2024

Balaan, C., Corley, M. J., Eulalio, T., Leite-Ahyo, K., Pang, A. P. S., Fang, R., Khadka, V. S., Maunakea, A. K., & Ward, M. A. (2019). Juvenile Shank3b deficient mice present with behavioral phenotype relevant to autism spectrum disorder. Behavioural Brain Research, 356, 137–147. 10.1016/j.bbr.2018.08.005

Balakrishnan, S., & Pearce, R.A. (2015) Spatiotemporal characteristics and pharmacological modulation of multiple gamma oscillations in the CA1 region of the hippocampus. Front Neural Circuits. 8:150. doi: 10.3389/fncir.2014.00150.

Barbaresi, P., Mensà, E., Bastioli, G., & Amoroso, S. (2017). Substance P NK1 receptor in the rat corpus callosum during postnatal development. Brain and Behavior, 7(6), e00713. 10.1002/brb3.713

Boggio, E. M., Ehlert, E. M., Lupori, L., Moloney, E. B., De Winter, F., Vander Kooi, C. W., Baroncelli, L., Mecollari, V., Blits, B., Fawcett, J. W., Verhaagen, J., & Pizzorusso, T. (2019). Inhibition of semaphorin3a promotes ocular dominance plasticity in the adult rat visual cortex. Molecular Neurobiology, 56(9), 5987–5997. 10.1007/s12035-019-1499-0

Borhegyi, Z., & Leranth, C. (1997). Substance P innervation of the rat hippocampal formation. Journal of Comparative Neurology, 384(1), 41–58. 10.1002/(SICI)1096-9861(19970721)384:1<41::AID-CNE3>3.0.CO;2-L

Brust, V., Schindler, P. M., & Lewejohann, L. (2015). Lifetime development of behavioural phenotype in the house mouse (Mus musculus). Frontiers in Zoology, 12 Suppl 1(Suppl 1), S17. 10.1186/1742-9994-12-S1-S17

Carr, M. F., Karlsson, M. P., & Frank, L. M. (2012). Transient slow gamma synchrony underlies hippocampal memory replay. Neuron, 75(4), 700–713.10.1016/j.neuron.2012.06.014

Carstens, K. E., Phillips, M. L., Pozzo-Miller, L., Weinberg, R. J., & Dudek, S. M. (2016). Perineuronal nets suppress plasticity of excitatory synapses on CA2 pyramidal neurons. The Journal of Neuroscience, 36(23), 6312–6320. 10.1523/JNEUROSCI.0245-16.2016

Carstens, K. E., Lustberg, D. J., Shaughnessy, E. K., McCann, K. E., Alexander, G. M., & Dudek, S. M. (2021). Perineuronal net degradation rescues CA2 plasticity in a mouse model of Rett syndrome. The Journal of Clinical Investigation, 131(16). 10.1172/JCI137221

Chen, S., He, L., Huang, A. J. Y., Boehringer, R., Robert, V., Wintzer, M. E., Polygalov, D., Weitemier, A. Z., Tao, Y., Gu, M., Middleton, S. J., Namiki, K., Hama, H., Therreau, L., Chevaleyre, V., Hioki, H., Miyawaki, A., Piskorowski, R. A., & McHugh, T. J. (2020). A hypothalamic novelty signal modulates hippocampal memory. Nature, 586(7828), 270–274. 10.1038/s41586-020-2771-1.

Chung, M., Imanaka, K., Huang, Z., Watarai, A., Wang, M.Y., Tao, K., Ejima, H., Aida, T., Feng. G., & Okuyama, T. (2024) Conditional knockout of Shank3 in the ventral CA1 by quantitative in vivo genome-editing impairs social memory in mice. Nat Commun,15(1), 4531. doi: 10.1038/s41467-024-48430-x.

Colgin, L.L. (2016). Rhythms of the hippocampal network. Nat Rev Neurosci. 17(4), 239–49. doi: 10.1038/nrn.2016.21.

Connolly, S., Anney, R., Gallagher, L., & Heron, E. A. (2017). A genome-wide investigation into parent-of-origin effects in autism spectrum disorder identifies previously associated genes including SHANK3. European Journal of Human Genetics, 25(2), 234–239. 10.1038/ejhg.2016.153

Cope, E. C., Zych, A. D., Katchur, N. J., Waters, R. C., Laham, B. J., Diethorn, E. J., Park, C. Y., Meara, W. R., & Gould, E. (2022). Atypical perineuronal nets in the CA2 region interfere with social memory in a mouse model of social dysfunction. Molecular Psychiatry, 27(8), 3520–3531. 10.1038/s41380-022-01559-0

Cope, E. C., Wang, S. H., Waters, R. C., Gore, I. R., Vasquez, B., Laham, B. J., & Gould, E. (2023). Activation of the CA2-ventral CA1 pathway reverses social discrimination dysfunction in Shank3B knockout mice. Nature Communications, 14(1), 1750. 10.1038/s41467-023-37248-8

Cui, Z., Gerfen, C. R., & Young 3rd, W. S. (2013). Hypothalamic and other connections with dorsal CA2 area of the mouse hippocampus. Journal of Comparative Neurology, 521(8), 1844–1866. 10.1002/cne.23263

Dasgupta, A., Baby, N., Krishna, K., Hakim, M., Wong, Y. P., Behnisch, T., Soong, T. W., & Sajikumar, S. (2017). Substance P induces plasticity and synaptic tagging/capture in rat hippocampal area CA2. Proceedings of the National Academy of Sciences, 114(41), E8741–E8749. 10.1073/pnas.1711267114

De Wit, J., De Winter, F., Klooster, J., & Verhaagen, J. (2005). Semaphorin 3A displays a punctate distribution on the surface of neuronal cells and interacts with proteoglycans in the extracellular matrix. Molecular and Cellular Neuroscience, 29(1), 40–55. 10.1016/j.mcn.2004.12.009

Dhamne, S. C., Silverman, J. L., Super, C. E., Lammers, S. H. T., Hameed, M. Q., Modi, M. E., Copping, N. A., Pride, M. C., Smith, D. G., Rotenberg, A., Crawley, J. N., & Sahin, M. (2017). Replicable in vivo physiological and behavioral phenotypes of the Shank3B null mutant mouse model of autism. Molecular Autism, 8(1), 26. 10.1186/s13229-017-0142-z

Diethorn, E. J., & Gould, E. (2023a). Postnatal development of hippocampal CA2 structure and function during the emergence of social recognition of peers. Hippocampus, 33(3), 208–222. 10.1002/hipo.23476

Diethorn, E. J., & Gould, E. (2023b). Development of the hippocampal CA2 region and the emergence of social recognition. Developmental Neurobiology, 83(5–6), 143–156. 10.1002/dneu.22919

Domínguez, S., Rey, C. C., Therreau, L., Fanton, A., Massotte, D., Verret, L., Piskorowski, R. A., & Chevaleyre, V. (2019). maturation of pnn and erbb4 signaling in area CA2 during adolescence underlies the emergence of pv interneuron plasticity and social memory. Cell Reports, 29(5), 1099–1112.e4. 10.1016/j.celrep.2019.09.044

Donoghue, T., Haller, M., Peterson, E. J., Varma, P., Sebastian, P., Gao, R., Noto, T., Lara, A. H., Wallis, J. D., Knight, R. T., Shestyuk, A., & Voytek, B. (2020). Parameterizing neural power spectra into periodic and aperiodic components. Nature Neuroscience, 23(12), 1655–1665. 10.1038/s41593-020-00744-x

Fawcett, J. W., Oohashi, T., & Pizzorusso, T. (2019). The roles of perineuronal nets and the perinodal extracellular matrix in neuronal function. Nature Reviews. Neuroscience, 20(8), 451–465. 10.1038/s41583-019-0196-3

Foscarin, S., Raha-Chowdhury, R., Fawcett, J. W., & Kwok, J. C. F. (2017). Brain ageing changes proteoglycan sulfation, rendering perineuronal nets more inhibitory. Aging, 9(6), 1607–1622. 10.18632/aging.101256

Gauthier, J., Spiegelman, D., Piton, A., Lafrenière, R. G., Laurent, S., St-Onge, J., Lapointe, L., Hamdan, F. F., Cossette, P., Mottron, L., Fombonne, E., Joober, R., Marineau, C., Drapeau, P., & Rouleau, G. A. (2009). Novel de novo SHANK3 mutation in autistic patients. American Journal of Medical Genetics. 150B(3), 421–424. 10.1002/ajmg.b.30822

Guillory, S. B., Baskett, V. Z., Grosman, H. E., McLaughlin, C. S., Isenstein, E. L., Wilkinson, E., Weissman, J., Britvan, B., Trelles, M. P., Halpern, D. B., Buxbaum, J. D., Siper, P. M., Wang, A. T., Kolevzon, A., & Foss-Feig, J. H. (2021). Social visual attentional engagement and memory in Phelan-McDermid syndrome and autism spectrum disorder: A pilot eye tracking study. Journal of Neurodevelopmental Disorders, 13(1), 58. 10.1186/s11689-021-09400-2

Halasy, K., Hajszan, T., Kovács, E. G., Lam, T.-T., & Leranth, C. (2004). Distribution and origin of vesicular glutamate transporter 2-immunoreactive fibers in the rat hippocampus. Hippocampus, 14(7), 908–918. 10.1002/hipo.20006

Hayani, H., Song, I., & Dityatev, A. (2018). Increased excitability and reduced excitatory synaptic input into fast-spiking CA2 interneurons after enzymatic attenuation of extracellular matrix. Frontiers in Cellular Neuroscience, 12. 10.3389/fncel.2018.00149

Hitti, F. L., & Siegelbaum, S. A. (2014). The hippocampal CA2 region is essential for social memory. Nature, 508(7494), 88–92. 10.1038/nature13028

Laham, B., Diethorn, E. J., & Gould, E. (2021). Newborn mice form lasting CA2-dependent memories of their mothers. Cell Reports, 34(4). 10.1016/j.celrep.2020.108668

Laham, B. J., Gore, I. R., Brown, C. J., & Gould, E. (2024). Adult-born granule cells modulate CA2 network activity during retrieval of developmental memories of the mother. eLife, 12, RP90600. 10.7554/eLife.90600

Li, M., Kinney, J. L., Jiang, Y.-Q., Lee, D. K., Wu, Q., Lee, D., Xiong, W.-C., & Sun, Q. (2023). Hypothalamic supramammillary nucleus selectively excites hippocampal CA3 interneurons to suppress CA3 pyramidal neuron activity. Journal of Neuroscience, 43(25), 4612–4624. 10.1523/JNEUROSCI.1910-22.2023

Liu, L., Zhang, Y., Men, S., Li, X., Hou, S.-T., & Ju, J. (2023). Elimination of perineuronal nets in CA1 disrupts GABA release and long-term contextual fear memory retention. Hippocampus, 33(7), 862–871. 10.1002/hipo.23503

Mehak, S. F., Shivakumar, A. B., Jijimon, F., Gupta, A., Pillai, V. G., & Gangadharan, G. (2025).Targeting CA2 perineuronal nets restores recognition memory and theta oscillations in aged mice. Aging Cell, 24(9), e70139. 10.1111/acel.70139

Mei, Y.-X., Chen, H.-S., Cao, Q.-L., Lv, W.-W., Liu, Y., Deng, S.-L., Zhou, Q., Zhang, Z.-J., Wang, F., Hu, Z.-L., & Chen, J.-G. (2025). Semaphorin 3A-mediated perineuronal nets formation incubates depressive-like behaviors in male mice via activating parvalbumin-expressing interneurons. Molecular Psychiatry, 1–17. 10.1038/s41380-025-03239-y

Mirzadeh, Z., Alonge, K. M., Cabrales, E., Herranz-Pérez, V., Scarlett, J. M., Brown, J. M., Hassouna, R., Matsen, M. E., Nguyen, H. T., Garcia-Verdugo, J. M., Zeltser, L. M., & Schwartz, M. W. (2019). Perineuronal net formation during the critical period for neuronal maturation in the hypothalamic arcuate nucleus. Nature Metabolism, 1(2), 212–221. 10.1038/s42255-018-0029-0

Morita, A., Yamashita, N., Sasaki, Y., Uchida, Y., Nakajima, O., Nakamura, F., Yagi, T., Taniguchi, M., Usui, H., Katoh-Semba, R., Takei, K., & Goshima, Y. (2006). Regulation of dendritic branching and spine maturation by semaphorin3a-fyn signaling. Journal of Neuroscience, 26(11), 2971–2980. 10.1523/JNEUROSCI.5453-05.2006

Okabe, S., Nagasawa, M., Kihara, T., Kato, M., Harada, T., Koshida, N., Mogi, K., & Kikusui, T. (2010). The effects of social experience and gonadal hormones on retrieving behavior of mice and their responses to pup ultrasonic vocalizations. Zoological Science, 27(10), 790–795. 10.2108/zsj.27.790

Okuyama, T., Kitamura, T., Roy, D.S., Itohara, S., & Tonegawa, S. (2016) Ventral CA1 neurons store social memory. Science. 353, 1536–1541.

Oliva, A., Fernández-Ruiz, A., Leroy, F., & Siegelbaum, S. A. (2020). Hippocampal CA2 sharp-wave ripples reactivate and promote social memory. Nature, 587(7833), Article 7833. 10.1038/s41586-020-2758-y

Phelan, K., & McDermid, H. E. (2011). The 22q13.3 Deletion Syndrome (Phelan-McDermid Syndrome). Molecular Syndromology, 2(3–5), 186–201. 10.1159/000334260

Pochinok, I., Stöber, T. M., Triesch, J., Chini, M., & Hanganu-Opatz, I. L. (2024). A developmental increase of inhibition promotes the emergence of hippocampal ripples. Nature Communications, 15(1), 738. 10.1038/s41467-024-44983-z

Rey, C. C., Robert, V., Bouisset, G., Loisy, M., Lopez, S., Cattaud, V., Lejards, C., Piskorowski, R. A., Rampon, C., Chevaleyre, V., & Verret, L. (2022). Altered inhibitory function in hippocampal CA2 contributes in social memory deficits in Alzheimer’s mouse model. iScience, 25(3). 10.1016/j.isci.2022.103895

Robert, V., Therreau, L., Chevaleyre, V., Lepicard, E., Viollet, C., Cognet, J., Huang, A. J., Boehringer, R., Polygalov, D., McHugh, T. J., & Piskorowski, R. A. (2021). Local circuit allowing hypothalamic control of hippocampal area CA2 activity and consequences for CA1. eLife, 10, e63352. 10.7554/eLife.63352

Stevenson, E. L., & Caldwell, H. K. (2014). Lesions to the CA2 region of the hippocampus impair social memory in mice. The European Journal of Neuroscience, 40(9), 3294–3301. 10.1111/ejn.12689

Tao, K., Chung, M., Watarai, A., Huang, Z., Wang, M.-Y., & Okuyama, T. (2022). Disrupted social memory ensembles in the ventral hippocampus underlie social amnesia in autism-associated Shank3 mutant mice. Molecular Psychiatry, 27(4), 2095–2105. 10.1038/s41380-021-01430-5

Tort, A. B. L., Komorowski, R., Eichenbaum, H., & Kopell, N. (2010). Measuring phase-amplitude coupling between neuronal oscillations of different frequencies. Journal of Neurophysiology, 104(2), 1195–1210. 10.1152/jn.00106.2010

Uesaka, N., Uchigashima, M., Mikuni, T., Nakazawa, T., Nakao, H., Hirai, H., Aiba, A., Watanabe, M., & Kano, M. (2014). Retrograde semaphorin signaling regulates synapse elimination in the developing mouse brain. Science, 344(6187), 1020–1023. 10.1126/science.1252514

Vertes, R.P. (2015) Major diencephalic inputs to the hippocampus: supramammillary nucleus and nucleus reuniens. Circuitry and function. Prog Brain Res. 219, 121–44. doi: 10.1016/bs.pbr.2015.03.008.

Vicente, A.F., Slézia, A., Ghestem, A., Bernard, C., Quilichini, P.P. (2020) In vivo characterization of neurophysiological diversity in the lateral supramammillary nucleus during hippocampal sharp-wave ripples of adult rats. Neuroscience. 435, 95–111. doi: 10.1016/j.neuroscience.2020.03.034.

Vo, T., Carulli, D., Ehlert, E. M. E., Kwok, J. C. F., Dick, G., Mecollari, V., Moloney, E. B., Neufeld, G., de Winter, F., Fawcett, J. W., & Verhaagen, J. (2013). The chemorepulsive axon guidance protein semaphorin3A is a constituent of perineuronal nets in the adult rodent brain. Molecular and Cellular Neurosciences, 56, 186–200. 10.1016/j.mcn.2013.04.009

Wang, D., & Fawcett, J. (2012). The perineuronal net and the control of CNS plasticity. Cell and Tissue Research, 349(1), 147–160. 10.1007/s00441-012-1375-y

Weigelt, S., Koldewyn, K., & Kanwisher, N. (2012). Face identity recognition in autism spectrum disorders: A review of behavioral studies. Neuroscience & Biobehavioral Reviews, 36(3), 1060–1084. 10.1016/j.neubiorev.2011.12.008

Wenzel, H. J., Cole, T. B., Born, D. E., Schwartzkroin, P. A., & Palmiter, R. D. (1997). Ultrastructural localization of zinc transporter-3 (ZnT-3) to synaptic vesicle membranes within mossy fiber boutons in the hippocampus of mouse and monkey. Proceedings of the National Academy of Sciences, 94(23), 12676–12681. 10.1073/pnas.94.23.12676

Yamashita, N., Morita, A., Uchida, Y., Nakamura, F., Usui, H., Ohshima, T., Taniguchi, M., Honnorat, J., Thomasset, N., Takei, K., Takahashi, T., Kolattukudy, P., & Goshima, Y. (2007). Regulation of spine development by semaphorin3a through cyclin-dependent kinase 5 phosphorylation of collapsin response mediator protein 1. The Journal of Neuroscience, 27(46), 12546–12554. 10.1523/JNEUROSCI.3463-07.2007

Zhu, N., Zhang, Y., Xiao, X., Wang, Y., Yang, J., Colgin, L. L., & Zheng, C. (2023). Hippocampal oscillatory dynamics in freely behaving rats during exploration of social and non-social stimuli. Cognitive Neurodynamics, 17(2), 411–429. 10.1007/s11571-022-09829-8

